# Learning gene networks under SNP perturbation using SNP and allele-specific expression data

**DOI:** 10.1101/2023.10.23.563661

**Authors:** Jun Ho Yoon, Seyoung Kim

**Author notes:** Most of the research in this manuscript was performed while Seyoung Kim was at CMU.

## Abstract

Allele-specific expression quantification from RNA-seq reads provides opportunities to study the control of gene regulatory networks by *cis*-acting and *trans*-acting genetic variants. Many existing methods performed a single-gene and single-SNP association analysis to identify expression quantitative trait loci (eQTLs), and placed the eQTLs against known gene networks for functional interpretation. Instead, we view eQTL data as a capture of the effects of perturbation of gene regulatory system by a large number of genetic variants and reconstruct a gene network perturbed by eQTLs. We introduce a statistical framework called CiTruss for simultaneously learning a gene network and *cis*-acting and *trans*-acting eQTLs that perturb this network, given population allele-specific expression and SNP data. CiTruss uses a multi-level conditional Gaussian graphical model to model *trans*-acting eQTLs perturbing the expression of both alleles in gene network at the top level and *cis*-acting eQTLs perturbing the expression of each allele at the bottom level. We derive a transformation of this model that allows efficient learning for large-scale human data. Our analysis of the GTEx and LG×SM advanced intercross line mouse data for multiple tissue types with CiTruss provides new insights into genetics of gene regulation. CiTruss revealed that gene networks consist of local subnetworks over proximally located genes and global subnetworks over genes scattered across genome, and that several aspects of gene regulation by eQTLs such as the impact of genetic diversity, pleiotropy, tissue-specific gene regulation, and local and long-range linkage disequilibrium among eQTLs can be explained through these local and global subnetworks.

## Introduction

To uncover unknown gene regulatory networks in an organism, it is necessary to perturb the biological system and to observe the effects of the perturbation. Naturally-occurring perturbation captured in expression and genetic variant data collected for a population has been considered as a powerful alternative to experimental perturbation, as it offers advantages of being less costly and having the potential for more meaningful discoveries that explain phenotypic variability found in nature ^1,2,3^. However, most of the existing methods were limited to expression quantitative trait locus (eQTL) mapping with total or allele-specific expression data, which at their core considered association between a single SNP and single gene expression. Here, we view eQTL data as a capture of the effects of perturbation of gene regulatory system by a large number of genetic variants and reconstruct a gene network perturbed by eQTLs.

Gene networks and eQTLs that perturb these networks inform each other, and thus should be identified simultaneously. Only the parts of networks that are not conserved but are perturbed by genetic variants can be reconstructed from eQTL data. The reconstructed network can then be used to isolate eQTLs with direct effects from those with indirect effects on the downstream genes. Identifying eQTLs first ^4,5,6,7^ and placing them on a known gene regulatory network has limitations, because the known network does not reflect the genetic background of samples specific to a given data and cannot help us disentangle pleiotropy.

Here, we introduce a statistical framework called CiTruss for reconstructing a gene network perturbed by *cis*-acting and *trans*-acting eQTLs given allele-specific expression and phased SNP data for a population of individuals. CiTruss uses a new class of probabilistic graphical model ^8^, called a multi-level conditional Gaussian graphical model, that extends the single-level conditional Gaussian graphical model, which was previously developed for learning gene networks perturbed by eQTLs ^9,10^, to additionally determine whether eQTLs are *cis*-acting or *trans*-acting. We describe two transformations of this multi-level model: a hybrid model to handle genes with unobserved allele-specific expression due to lack of heterozygous loci in the coding regions and a sum-difference model to allow for efficient learning with large-scale human data. While the estimated multi-level model contains eQTLs with direct effects, eQTLs with indirect effects from pleiotropy on gene network are obtained by performing inference on this model. In the special case of gene network with no edges, our multi-level model reduces to a scaled multivariate linear regression model.

CiTruss can be used as a standalone method or can be paired with an existing statistical method for eQTL mapping ^11,12,7,6^. As a partner method, CiTruss takes eQTLs and the genes affected by eQTLs, called eGenes, found by the existing method as well as the original eQTL data, and outputs a gene network and eQTLs with direct effects acting in *cis* or *trans*, after eliminating eQTLs with indirect effects. We show that CiTruss combined with Matrix eQTL is an effective and efficient approach to obtaining gene networks and eQTLs with a measure of statistical significance.

Applying CiTruss to data from the unrelated individuals of the GTEx Project ^13^ and from the LG×SM advanced intercross line (AIL) mice in generations 50 to 56^14^ shows that the difference in genetic diversity between the two populations leads to different gene networks. The gene networks for mice were more compact and focused due to fewer genetic variants that affect the difference in a narrower set of behavioral traits between the two founder strains. The two populations also shared several similarities in gene networks and eQTLs. First, gene networks consisted of local subnetworks over proximally located genes or global subnetworks over genes scattered across genome. Second, the pleiotropic effects of eQTLs were organized around these local and global subnetworks, which CiTruss disentangled into *cis*-acting and *trans*-acting eQTLs with direct effects. Third, gene networks from the two GTEx tissue types, whole blood and muscle skeletal, and the three mouse tissue types, prefrontal cortex (PFC), striatum (STR), and hippocampus (HIP), had global subnetworks with tissue-specific genes and eQTLs, but local subnetworks that were often not tissue-specific. Long-range linkage disequilibrium in eQTLs for global subnetworks was found only in the GTEx data, and thus, may have played a role in shaping tissue-specific gene regulation during evolution. Our results substantially elaborate the previous reports on the pleiotropic effects of eQTLs on clusters of locally co-expressed genes ^15^, a hierarchy in gene expression and eQTL patterns across tissue types ^16^, and long-range linkage disequilibrium that recent studies have begun to notice in genome-wide association studies ^17,18,19^.

## Results

### Method overview

CiTruss is a statistical framework that consists of three components: model, learning, and inference. For model, CiTruss has a multi-level conditional Gaussian graphical model with two sets of parameters, one for a gene network and the other for perturbation of this network by *cis*-acting and *trans*-acting eQTLs (Fig. 1(a)). In this multi-level model, each allele of *cis*-acting eQTL affects the expression of the gene allele on the same haplotype and each *trans*-acting eQTL affects the expression of both gene alleles in the gene network. When allele-specific expression cannot be measured because of the absence of heterozygous loci in the gene coding region, the multi-level model collapses to a hybrid model that consists of multi-level and single-level models, each for genes with and without observed allele-specific expression (Figs. 1(b)-(c)). We show that both the multi-level model and hybrid model can be transformed to a sum-difference model that factorizes to two single-level models: a sum model for the total expression of both alleles perturbed by either *cis*-acting or *trans*-acting eQTLs and a difference model for the differential expression of two alleles perturbed by *cis*-acting eQTLs (Figs. 1(d)-(f)). The sum-difference model is used for efficient learning, where each of the two single-level models is estimated with the efficient learning algorithm Mega-sCGGM that was previously developed for single-level models ^10^. With inference, CiTruss infers the indirect effects of eQTLs, as *cis*-acting or *trans*-acting eQTLs with direct effects in the multi-level model become *trans*-acting eQTLs with indirect effects on the other genes in the network (Fig. S1).

**Figure 1.**
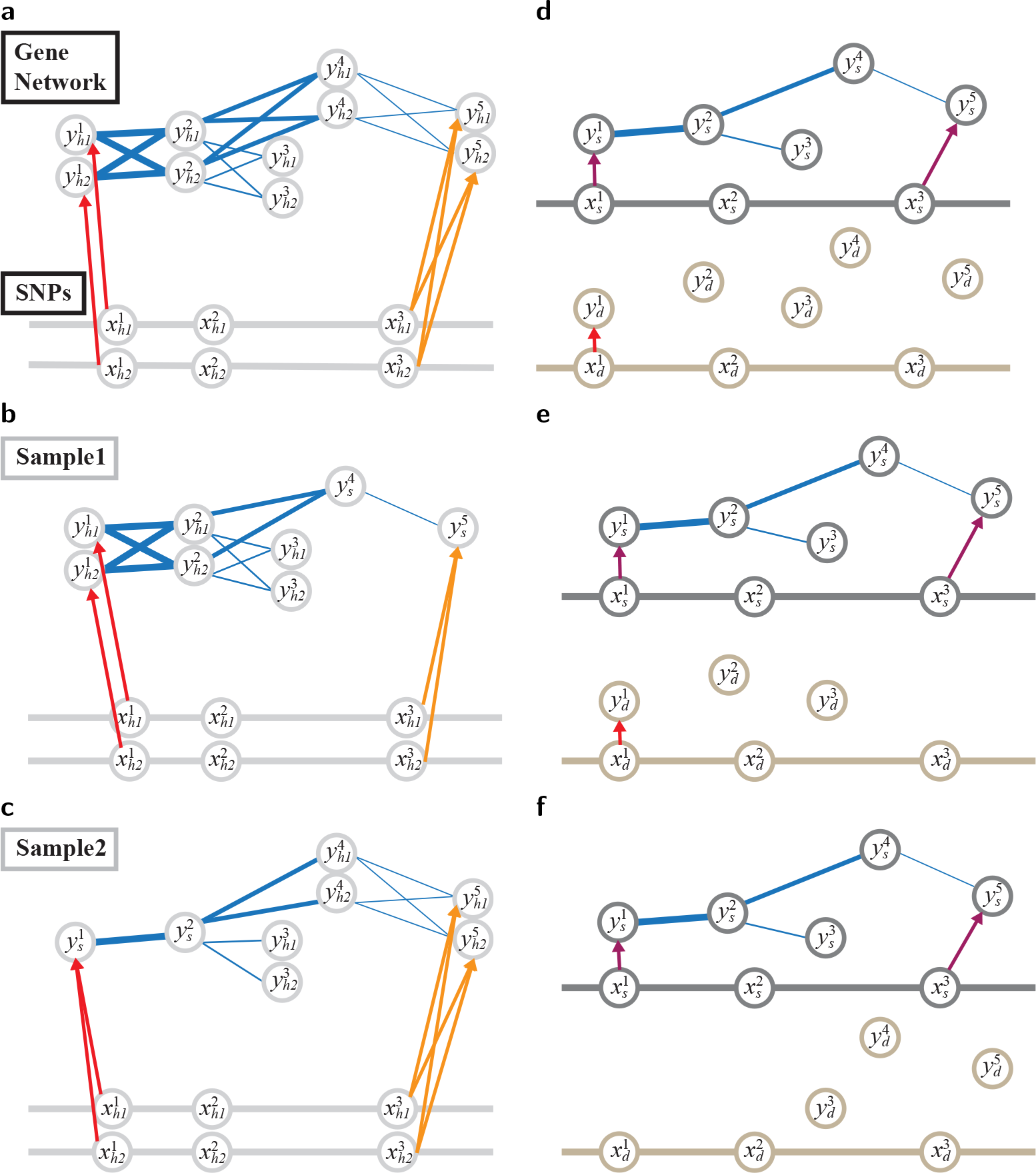
Overview of CiTruss. (a) Multi-level conditional Gaussian graphical model in CiTruss for a gene network perturbed by *cis*-acting and *trans*-acting eQTLs. At *cis*-acting eQTL (red edges), the SNP allele on each haplotype (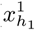 or 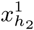 of SNP 1) affects the expression of the gene allele (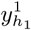 or 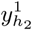 of gene 1) on the same haplotype. At *trans*-acting eQTL (orange edges), the SNP alleles on both haplotypes (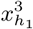 and 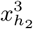) affect the expression of both gene alleles (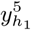 and 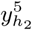). In gene network (blue edges), the interaction between gene *i* and gene *j* is represented as four edges between the two alleles (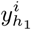 and 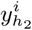) of gene *i* and the two alleles (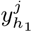 and 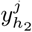) of gene *j*. (b) Hybrid model for sample 1. For genes 4 and 5 with unobserved allele-specific expression, the two nodes for two alleles collapse into a single node. (c) Hybrid model for sample 2 with unobserved allele-specific expression for gene 1. (d) Sum-difference model derived from the multi-level model in Panel (a). In the sum model (top), the sum of the expression of two alleles 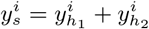 of gene *i* is perturbed by the genotype 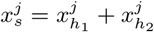 of SNP *j*. In the difference model (bottom), the differential expression of the two alleles 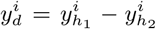 of gene *i* is affected by the difference of SNP alleles 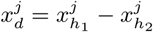 for SNP *j*. (e) Sum-difference model derived from the hybrid model in Panel (b). (f) Sum-difference model derived from the hybrid model in Panel (c).

### Simulation

Using simulated data with known ground truth, we benchmarked CiTruss against TReCASE ^12^, RASQUAL ^6^, WASP ^7^, and Matrix eQTL ^11^. We used small datasets, each with 100 SNPs, 40 genes, and 200 samples, because TReCASE and WASP required several hours of computation time even for a dataset of this size. CiTruss outperformed all methods on the accuracy of eQTLs (Fig. S2). As the frequency of heterozygous genotypes and the frequency of observed allele-specific expression increased, the information necessary to detect *cis*-acting eQTLs increased, and the accuracy of *cis*-acting eQTLs improved for all methods, but the accuracy of *trans*-acting eQTLs remained unaffected. Matrix eQTL almost always detected true *cis*-acting and *trans*-acting eQTLs as eQTLs with higher accuracy than any of the existing methods for allele-specific eQTL mapping. In re-analysis of the eQTLs (FDR < 0.05) from the existing methods with CiTruss, CiTruss eliminated many of the indirect eQTLs from the existing methods, while improving power and reducing false discovery rate (Figs. S3 and S4). On the accuracy of gene networks, CiTruss alone performed similarly or slightly better than CiTruss combined with an existing method (Fig. S5).

### Computation time

We compared the computation time of CiTruss and the existing methods, using the simulated, GTEx, and LG×SM AIL mouse data (Fig. 2). For the simulated dataset, CiTruss took two seconds, which was two, three, and four orders of magnitude faster than RASQUAL, TReCASE, and WASP, respectively. On the GTEx data from whole blood with 1,175,808 SNPs, 10,636 genes, and 670 samples and muscle skeletal with 1,172,754 SNPs, 12,475 genes, and 706 samples, only CiTruss and Matrix eQTL were sufficiently fast, each requiring around three hours on a 128 core machine. CiTruss was nearly as fast as Matrix eQTL on the GTEx data, as CiTruss takes advantage of the high level of sparsity in the parameters. On the mouse data from PFC with 30,066 SNPs, 8,030 genes, and 208 samples, STR with 29,139 SNPs, 8,423 genes, and 189 samples, and HIP with 30,742 SNPs, 8,298 genes, and 239 samples, we applied TReCASE to only those SNP-gene pairs with less than 1Mb apart due to high computational cost, and compared the computation time per SNP-gene pair. CiTruss was five orders of magnitude faster than TReCASE, and required around four hours for all SNPs and genes on a 8 core machine. Applying CiTruss to the results from Matrix eQTL required nearly the same amount of time as running CiTruss alone.

**Figure 2.**
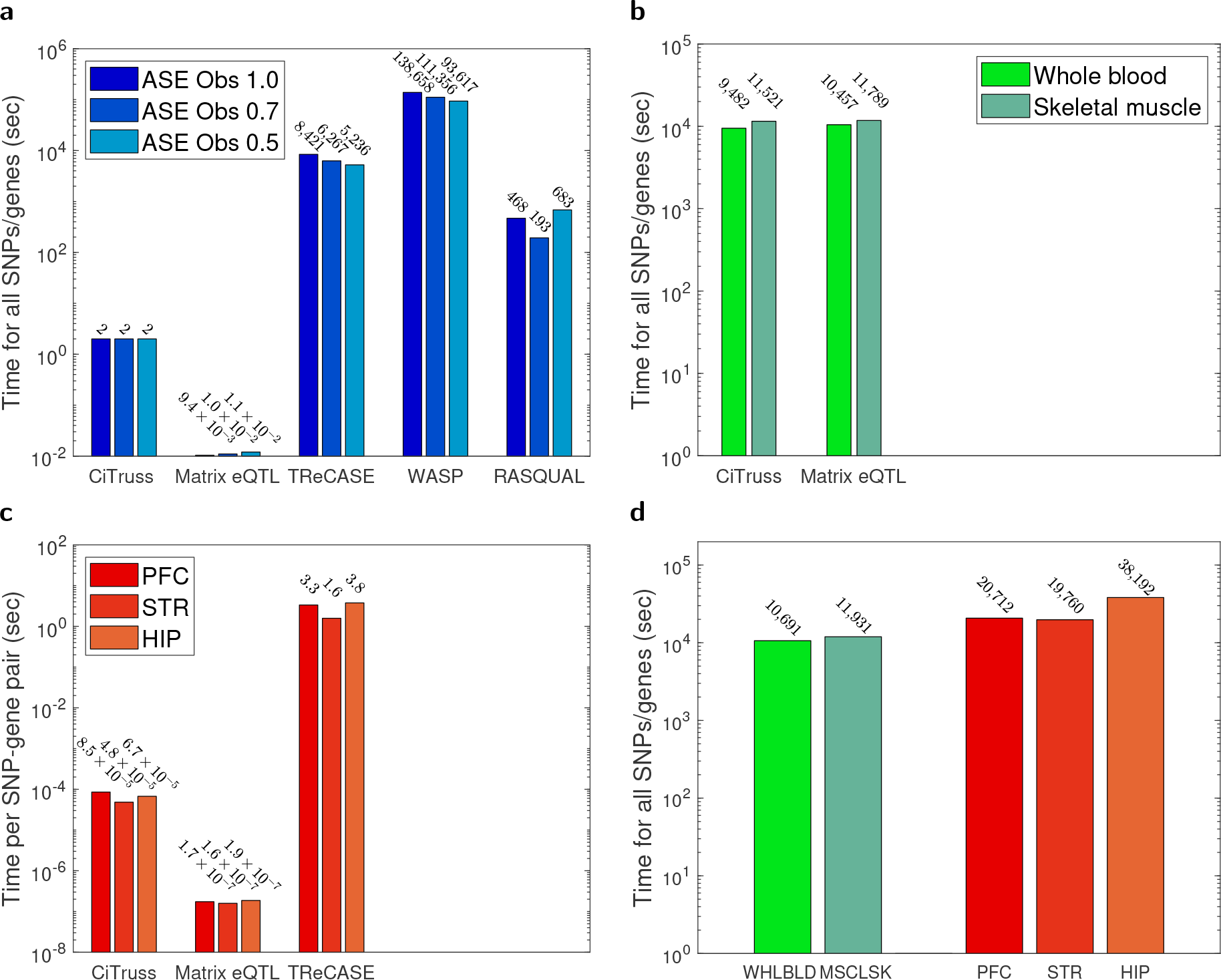
Computation time of CiTruss and other methods. (a) Simulated data. The frequency of heterozygous genotypes 0.25 was used in simulation. Time averaged over five simulated datasets is shown. (b) GTEx data with all SNPs and all genes. (c) LG×SM AIL mouse data. Time per SNP-gene pair is shown. (d) Computation time of CiTruss in re-analysis of Matrix eQTL results from the GTEx and LG×SM AIL mouse data.

### GTEx gene networks with local and global subnetworks

We applied CiTruss to the GTEx data for whole blood and muscle skeletal. Matrix eQTL was the only existing method that was sufficiently efficient for analysis of all SNPs and genes, and CiTruss combined with Matrix eQTL and as a standalone method led to qualitatively similar gene networks and eQTLs, indicating that Matrix eQTL is an effective tool for screening for eQTLs with both direct and indirect effects prior to applying CiTruss. Below, we focus on the results from applying CiTruss to eQTLs from Matrix eQTL (FDR < 0.05), and examine the gene network and *cis*-acting and *trans*-acting eQTLs as well as undetermined eQTLs, for which we could not determine whether they are acting in *cis* or *trans* due to a lack of samples with heterozygous genotypes.

In both tissue types, the gene networks from CiTruss consisted of isolated genes with no edges and connected genes in either one of many small local subnetworks or one relatively large global subnetwork (Fig. 3 whole blood; Fig. S6 muscle skeletal). The majority of the genes (74.6% whole blood; 68.0% muscle skeletal) were connected genes. The majority of these connected genes (76.8% whole blood; 93.0% muscle skeletal) belonged to one of the local subnetworks, each consisting of 2 to 249 genes located nearby on genome. The rest were in a global subnetwork over genes scattered across genome that could be seen when genes were clustered based on edges. A small portion of the connected genes (16.3% whole blood; 2.4% muscle skeletal) served as bridges between the local and global subnetworks. While co-expression of local genes in the GTEx data has been observed previously ^15^, CiTruss additionally learned a network over the co-expressed local genes and revealed global subnetworks that were well separated from the local subnetworks.

**Figure 3.**
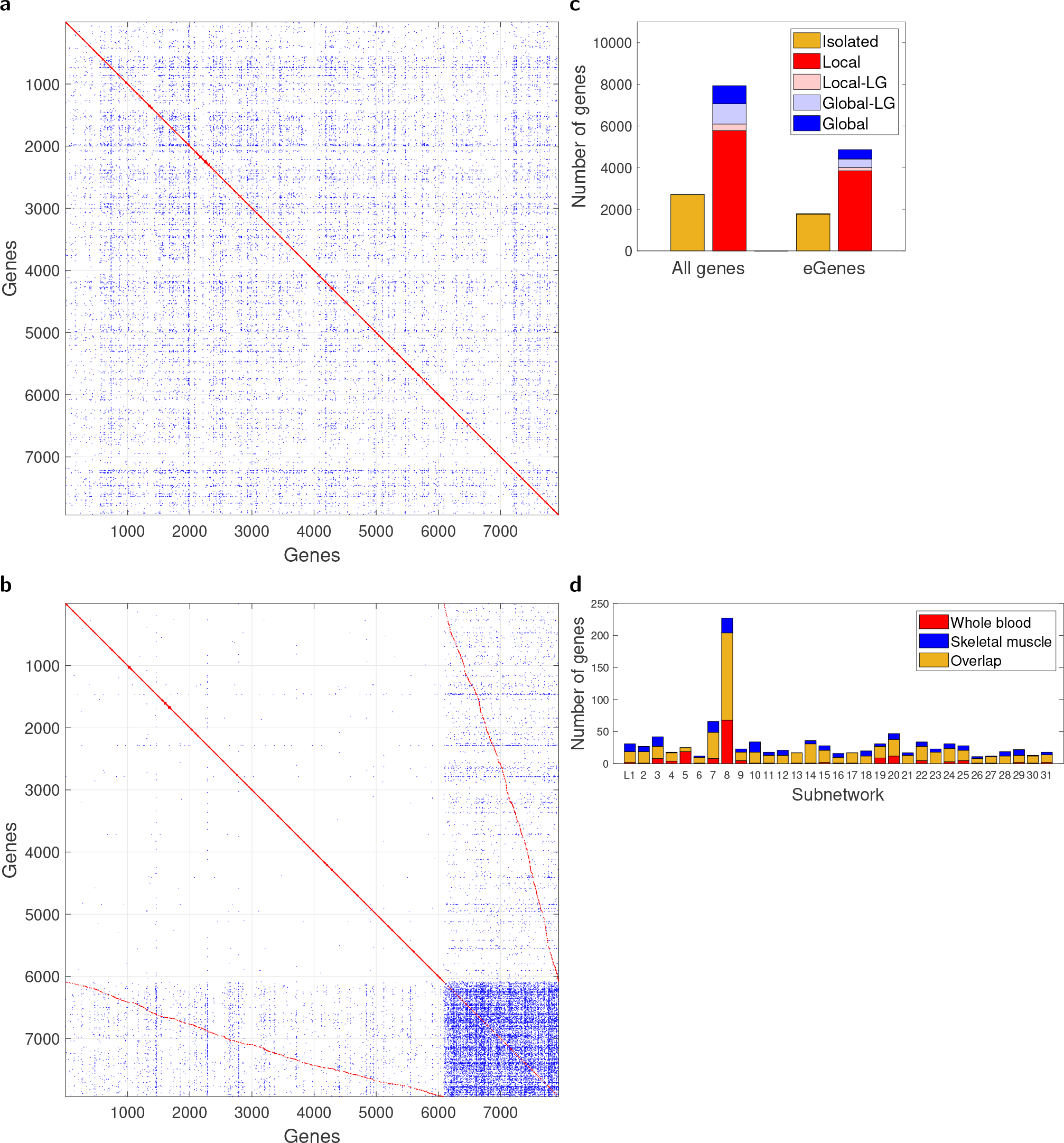
CiTruss gene network from whole blood GTEx data. (a) Gene network when genes are ordered according to their genome locations. If genes *i* and *j* are connected with an edge, (*i, j*) is marked with a dot (red for gene pairs with local connections defined as less than 50 genes apart; blue for pairs with non-local connections). Isolated genes with no edges are not shown. (b) Gene network when genes are reordered after clustering the network in Panel (a). The upper-left block shows genes and edges in the local subnetworks, and the lower-right block shows those in global subnetwork. (c) The number of genes and eGenes in different categories of network connectivity. Local-LG (and Global-LG): genes in the local subnetworks (and global subnetwork) that are connected to genes in the global subnetwork (and local subnetworks) with edges in the off-diagonal block in Panel (b). (d) Overlap in genes in the largest local subnetworks L1-L31 (Table S1) between whole blood and muscle skeletal.

Across tissue types, almost all of the large local subnetworks were not tissue-specific, but the global subnetworks were tissue-specific. The largest 31 local subnetworks L1–L31 had a large overlap in genes between the two tissue types (Fig. 3(d); Table S1), whereas the global subnetworks Gwb for whole blood and Gms for muscle skeletal had a small overlap in genes (128 between 1,848 in Gwb and 588 in Gms). For 22 of the 31 local subnetworks, the enriched gene ontology (GO) terms were not tissue specific, but the global subnetworks were enriched for genes with tissue-specific GO terms (Table 1).

**Table 1:** GO terms enriched in CiTruss subnetworks from GTEx data.

|  | Whole blood |  |  | Muscle skeletal |  |  | GO |  |
| --- | --- | --- | --- | --- | --- | --- | --- | --- |
|  | Net | Over | *p*-val | Net | Over | *p*-val |  |  |
| ID | Size | lap | [FDR] | Size | lap | [FDR] | Term | Size |
| L1 | 19 | 2 | 1.8×10<sup>-3</sup> | 29 | 2 | 2.7×10<sup>-3</sup> | BP Tumor necrosis factor-med. signaling pathway | 57 |
| L2 | 19 | 10 | 1.6×10<sup>-3</sup> | 26 | 17 | 9.4×10<sup>-5</sup> | BP Cellular metabolic process | 5828 |
| L3 | 27 | 2 | 9.3×10<sup>-5</sup> | 34 | 2 | 2.3×10<sup>-4</sup> | BP Golgi vesicle budding | 10 |
| L4 | 17 | 2 | 7.1×10<sup>-5</sup> | 14 | 2 | 2.8×10<sup>-5</sup> | BP Neg. reg. of thymocyte apoptotic process<sup>†</sup> | 10 |
| L5 | 25 | 23 | 1.0×10<sup>-49</sup> | 6 | 6 | 1.1×10<sup>-13</sup> | BP Immunoglobulin production<sup>†</sup> | 141 |
| L6 | 10 | 1 | 1.8×10<sup>-3</sup> | 11 | 1 | 1.6×10<sup>-3</sup> | BP Lipid droplet fusion | 3 |
| L7 | 49 | 3 | 1.8×10<sup>-6</sup> | 58 | 4 | 3.1×10<sup>-8</sup> | MF Hyalurononglucosaminidase activity | 8 |
| L8 | 204 | 28 | 2.8×10<sup>-36</sup> | 159 | 22 | 1.3×10<sup>-29</sup> | BP Presentation of peptide antigen<sup>†</sup> | 71 |
| L9 | 18 | 2 | 9.5×10<sup>-4</sup> | 18 | 2 | 6.2×10<sup>-4</sup> | BP Glycosyl compound metabolic process | 85 |
| L10 | 18 | 1 | 2.9×10<sup>-3</sup> | 33 | 1 | 5.6×10<sup>-3</sup> | BP Reg. of smooth muscle cell-matrix adhesion* | 4 |
| L11 | 13 | 2 | 6.7×10<sup>-5</sup> | 17 | 2 | 8.7×10<sup>-5</sup> | BP Pos. reg. of ruffle assembly | 15 |
| L12 | 13 | 1 | 1.9×10<sup>-3</sup> | 21 | 1 | 3.1×10<sup>-3</sup> | BP Neg. reg. of protein K48-linked deubiquitination | 2 |
| L13 | 17 | 2 | 1.5×10<sup>-3</sup> | 16 | 2 | 1.3×10<sup>-3</sup> | BP Reg. of osteoclast differentiation | 72 |
| L14 | 31 | 2 | 7.8×10<sup>-5</sup> | 35 | 2 | 8.9×10<sup>-5</sup> | BP Neg. reg. of relaxation of muscle* | 7 |
| L15 | 21 | 1 | 1.3×10<sup>-4</sup> | 26 | 2 | 1.4×10<sup>-4</sup> | BP Muscle filament sliding* | 11 |
| L16 | 10 | 1 | 2.6×10<sup>-3</sup> | 15 | 2 | 1.0×10<sup>-5</sup> | BP Reg. of protein localization to microtubule* | 5 |
| L17 | 17 | 1 | 1.3×10<sup>-3</sup> | 16 | 1 | 1.2×10<sup>-3</sup> | BP Pos. reg. chaperone-med. protein complex assembly | 1 |
| L18 | 12 | 1 | 1.0×10<sup>-2</sup> | 20 | 2 | 1.2×10<sup>-4</sup> | BP Resolution of meiotic recombination intermediates | 17 |
| L19 | 27 | 1 | 2.6×10<sup>-3</sup> | 22 | 1 | 2.1×10<sup>-3</sup> | BP Neg. reg. of macropinocytosis | 1 |
| L20 | 38 | 2 | 4.4×10<sup>-4</sup> | 35 | 2 | 3.7×10<sup>-4</sup> | MF Reg. of gene expression by genomic imprinting | 17 |
| L21 | 13 | 1 | 2.4×10<sup>-3</sup> | 17 | 1 | 3.4×10<sup>-3</sup> | BP Traversing start control point of mitotic cell cycle | 4 |
| L22 | 27 | 4 | 7.2×10<sup>-4</sup> | 29 | 4 | 1.2×10<sup>-3</sup> | BP Activation of immune response<sup>†</sup> | 370 |
| L23 | 18 | 3 | 1.8×10<sup>-4</sup> | 23 | 3 | 4.1×10<sup>-4</sup> | BP Regulation of DNA recombination | 135 |
| L24 | 24 | 3 | 8.7×10<sup>-7</sup> | 28 | 3 | 1.9×10<sup>-6</sup> | CC Gamma-tubulin complex* | 16 |
| L25 | 21 | 3 | 2.5×10<sup>-4</sup> | 23 | 2 | 3.9×10<sup>-3</sup> | BP Protein acylation | 160 |
| L26 | 8 | 1 | 8.7×10<sup>-4</sup> | 10 | 1 | 8.7×10<sup>-4</sup> | BP Very long-chain fatty acid beta-oxidation | 1 |
| L27 | 11 | 1 | 1.9×10<sup>-2</sup> | 11 | 2 | 2.0×10<sup>-4</sup> | BP Release of sequestered calcium ion into cytosol | 63 |
| L28 | 12 | 11 | 1.2×10<sup>-10</sup> | 19 | 15 | 4.7×10<sup>-13</sup> | BP Reg. of transcription by RNA polymerase II | 2594 |
| L29 | 13 | 1 | 1.6×10<sup>-3</sup> | 20 | 1 | 2.9×10<sup>-3</sup> | BP Acetyl-CoA biosynthetic process from acetate | 2 |
| L30 | 13 | 1 | 1.2×10<sup>-3</sup> | 12 | 1 | 1.0×10<sup>-3</sup> | BP Nuclear migration along microfilament | 2 |
| L31 | 14 | 1 | 9.7×10<sup>-4</sup> | 16 | 1 | 1.4×10<sup>-3</sup> | BP Peptidyl-lysine butyrylation | 1 |
| Gwb | 1847 | 357 | [1.4×10<sup>-27</sup>] |  |  |  | BP Immune system process<sup>†</sup> | 2256 |
|  |  | 277 | [7.3×10<sup>-25</sup>] |  |  |  | BP Immune response<sup>†</sup> | 1625 |
|  |  | 135 | [2.1×10<sup>-16</sup>] |  |  |  | BP Adaptive immune response<sup>†</sup> | 654 |
|  |  | 226 | [1.4×10<sup>-14</sup>] |  |  |  | BP Reg. of immune system process<sup>†</sup> | 1480 |
|  |  | 161 | [2.2×10<sup>-12</sup>] |  |  |  | BP Pos. reg. of immune system process<sup>†</sup> | 972 |
|  |  | 140 | [9.6×10<sup>-10</sup>] |  |  |  | BP Reg. of immune response<sup>†</sup> | 876 |
|  |  | 105 | [3.6×10<sup>-9</sup>] |  |  |  | BP Reg. of leukocyte activation<sup>†</sup> | 595 |
|  |  | 92 | [1.3×10<sup>-8</sup>] |  |  |  | BP Reg. of lymphocyte activation<sup>†</sup> | 503 |
|  |  | 34 | [4.2×10<sup>-5</sup>] |  |  |  | MF Antigen binding<sup>†</sup> | 124 |
|  |  | 49 | [2.3×10<sup>-9</sup>] |  |  |  | CC Immunoglobulin complex<sup>†</sup> | 169 |
|  |  | 31 | [1.0×10<sup>-3</sup>] |  |  |  | CC T cell receptor complex<sup>†</sup> | 149 |
| Gms |  |  |  | 590 | 47 | [2.7×10<sup>-3</sup>] | CC Supramolecular polymer* | 1050 |
|  |  |  |  |  | 45 | [4.3×10<sup>-3</sup>] | CC Supramolecular fiber* | 1042 |
|  |  |  |  |  | 11 | [9.4×10<sup>-3</sup>] | CC Keratin filament* | 102 |
|  |  |  |  |  | 52 | [2.5×10<sup>-2</sup>] | CC Supramolecular complex* | 1414 |
|  |  |  |  |  | 15 | [2.8×10<sup>-2</sup>] | CC Sarcomere* | 217 |
|  |  |  |  |  | 15 | [4.8×10<sup>-2</sup>] | CC Myofibril* | 238 |

<sup>†</sup>Blood GO terms
\*Muscle GO terms

### GTEx Type I and II blocks of eQTLs in local and long-range linkage disequilibrium for local and global subnetworks

We placed the hotspot eQTLs with more than 10 eGenes from Matrix eQTL against the CiTruss gene network to gain insights into the functional mechanisms underlying the pleiotropy. These hotspot eQTLs appeared in one of two types of eQTL blocks: Type I block for eQTLs in linkage disequilibrium that affect primarily a local subnetwork and a few genes in the global subnetwork; and Type II block for eQTLs in long-range linkage disequilibrium that affect primarily the global subnetwork and a few genes in local subnetworks (Fig. 4 whole blood; Fig. S7 muscle skeletal).

**Figure 4.**
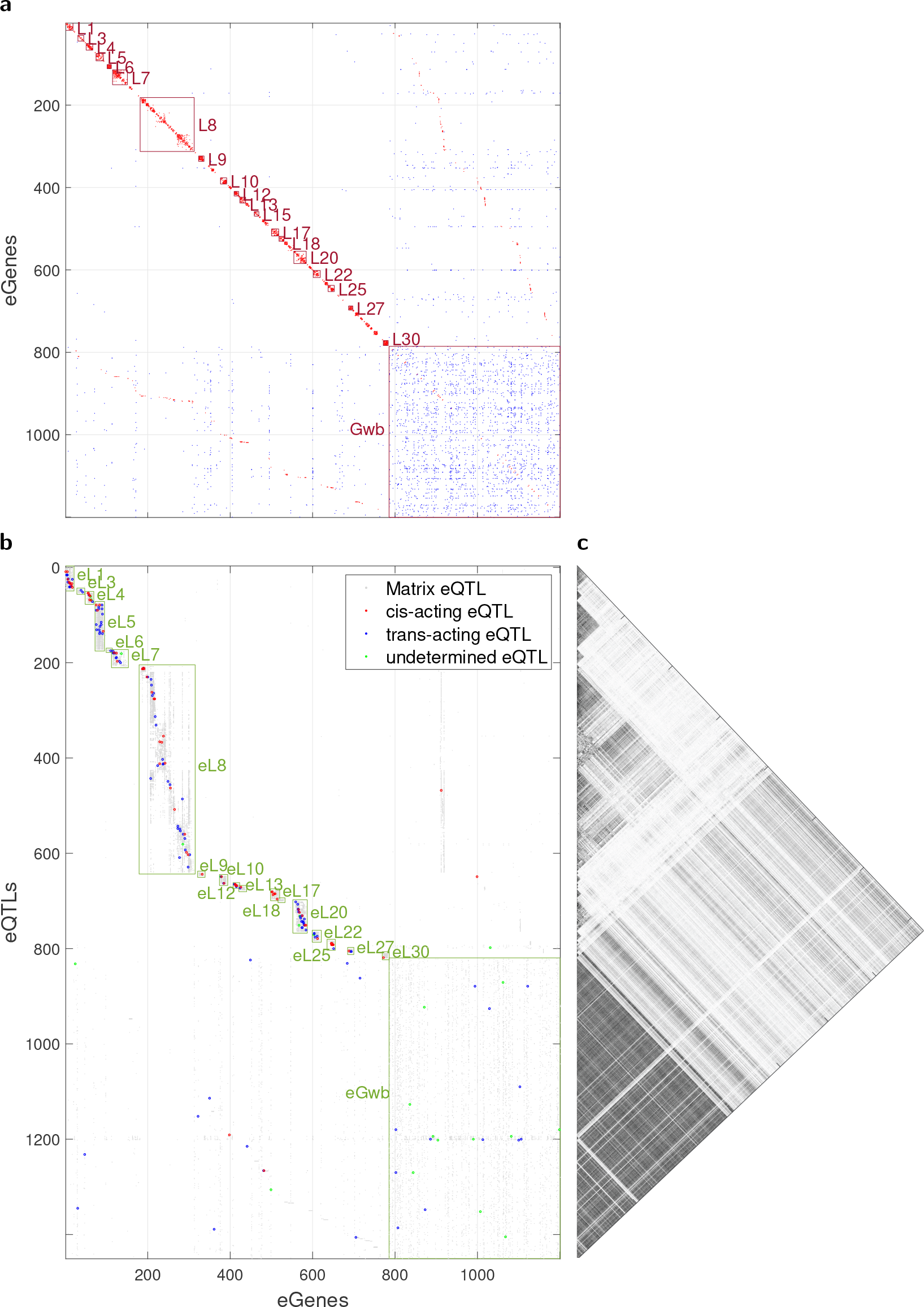
Matrix eQTL hotspots against CiTruss gene networks perturbed by eQTLs in whole blood GTEx data. (a) CiTruss gene network over eGenes of hotspot eQTLs (FDR*<*0.05, 10 or more eGenes) from Matrix eQTL. The local subnetworks (L1, etc.) and global subnetwork (Gwb) are shown. (b) Type I and Type II eQTL hotspot blocks, and CiTruss eQTLs with direct effects. Type I blocks are found from hotspot regions (eL1, etc.) and local subnetworks (L1, etc.). The single Type II block is found from many hotspot eQTLs scattered across genome (eGwb) and the global subnetwork (Gwb). See Table S1 for locations of hotspot regions and local subnetworks. (c) *r*^2^ for linkage disequilibrium in the Type I blocks and long-range linkage disequilibrium in the Type II block in Panel (b).

Most of the hotspot eQTLs (820 out of 1,451 whole blood; 991 out of 1,596 muscle skeletal) appeared in one of Type I blocks, each formed by one of hotspots eL1-eL31 and local subnetworks L1–L31. In each block, the position of the hotspot coincided with that of the corresponding local subnetwork (Table S1), which indicates these eQTLs are proximal to their eGenes. Although all local subnetworks L1–L31 were shared between the two tissue types, only 13 out of the 31 local subnetworks had hotspot eQTLs in both tissue types and thus shared the corresponding Type I block. The other 18 local subnetworks had hotspot eQTLs in only one of the two tissue types, several of which were enriched for genes with tissue-specific GO terms (Tables 1 and S1), suggesting that some local subnetworks may be involved in tissue-specific gene regulation through different genetic control for different tissue types.

The rest of the hotspot eQTLs appeared in a single Type II block for the global subnetwork. These eQTLs were in long-range linkage disequilibrium (eGwb, max *r*^2^ = 0.93, median *r*^2^ = 0.52, whole blood; eGms, max *r*^2^ = 0.93, median *r*^2^ = 0.52, muscle skeletal). While Type I blocks have been previously reported as eQTLs for co-expressed local genes ^15^, our analysis revealed previously unknown Type II blocks. Long-range linkage disequilibrium among SNPs has been known to arise from epistatic selection or population structure, and to be implicated in complex diseases ^17,18,19^, but little has been known about their roles in gene regulation. Our results suggest that long-range linkage disequilibrium may have played a role in shaping tissue-specific gene regulation in global subnetworks.

### GTEx eQTLs with direct effects

CiTruss selected a small number of the eQTL-eGene pairs from Matrix eQTL (10,308 out of 505,630 in whole blood; 12,562 out of 578,289 in muscle skeletal) as eQTLs with direct effects (Fig. 5(a)-(b) whole blood; Fig. S8(a)-(b) muscle skeletal). The majority of the *cis*-acting and *trans*-acting eQTLs were found in the local subnetworks, and a far smaller fraction in the global subnetwork. Although undetermined eQTLs were only a small fraction of all eQTLs (14.2% whole blood; 16.5% muscle skeletal), they represented the largest proportion of the eQTLs for the global subnetworks (49.3% whole blood; 79.1% muscle skeletal). Consistent with previous studies, CiTruss found nearly all *cis*-acting eQTLs proximal to their eGenes within 10Mb (Fig. S9), even though it was not constrained to do so and instead considered all eQTL-eGene pairs on the same chromosome from Matrix eQTL as candidate *cis*-acting eQTL-eGene pairs. However, different from previous reports, a large fraction of *trans*-acting eQTLs (93.6% whole blood; 96.6% muscle skeletal) were also located near their eGenes within 10Mb.

**Figure 5.**
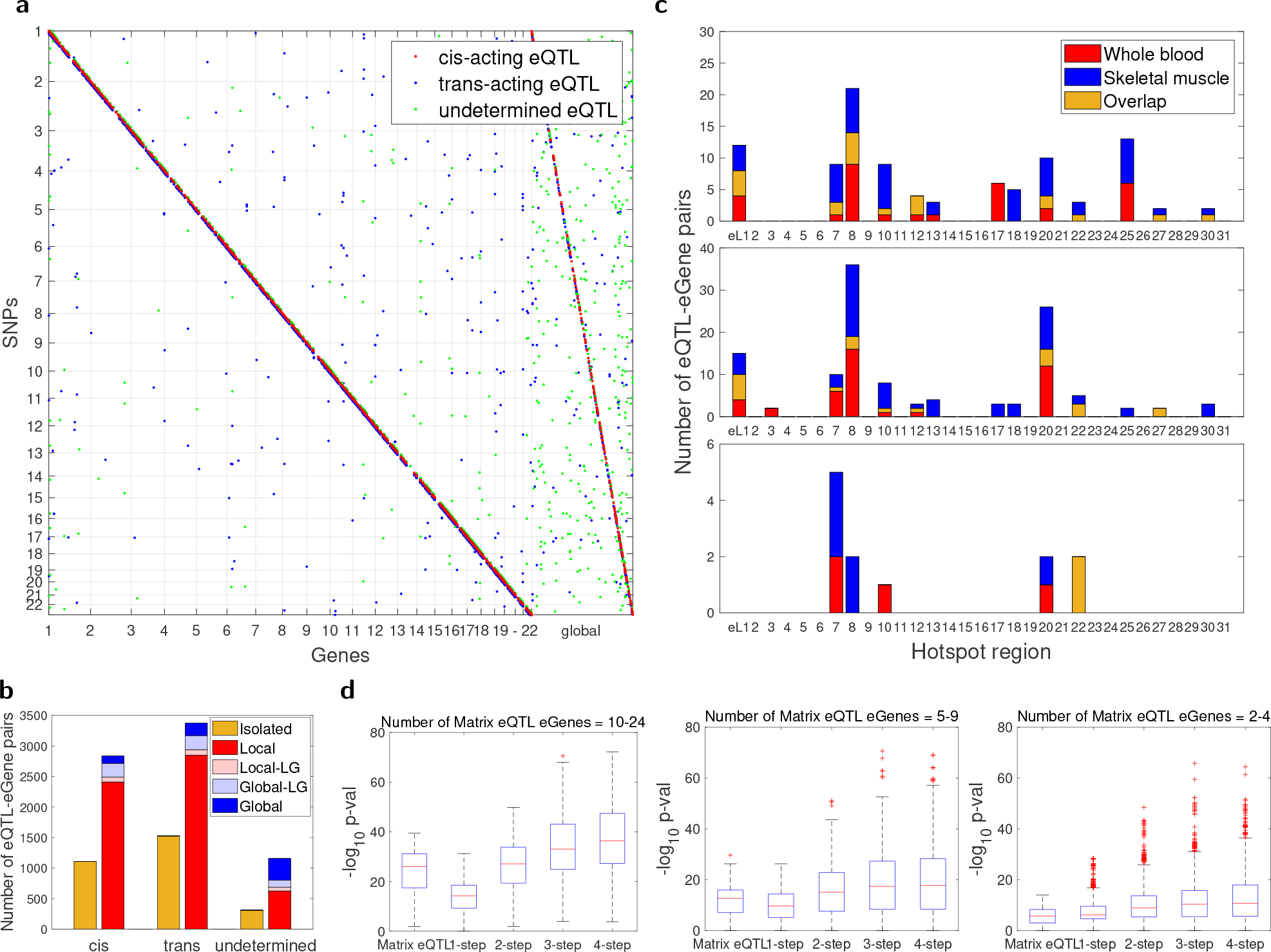
CiTruss eQTLs from whole blood GTEx data. (a) Locations of eQTLs vs eGenes, after grouping genes according to local and global subnetworks. (b) The number of eQTL-eGene pairs in different categories of network connectivity. (c) The overlap of eQTLs for local subnetworks between whole blood and muscle skeletal. *Cis*-acting eQTLs (top), *trans*-acting eQTLs (middle), and undetermined eQTLs (bottom). (d) Overlap between eQTL-eGene pairs and enhancer-target pairs. Each boxplot shows the distribution of hypergeometric test *p*-values over eQTL-eGene pairs. For CiTruss, the pair of each *cis*-acting eQTL and the set of the corresponding eGene and its 1, 2, 3, and 4 step network neighbors was used. For Matrix eQTL, the eQTLs selected as *cis*-acting by CiTruss were used.

CiTruss disentangled Type I eQTL blocks to *cis*-acting and *trans*-acting eQTLs for the local subnetworks and *trans*-acting eQTLs for the global subnetwork, and Type II blocks to primarily *trans*-acting and undetermined eQTLs and few *cis*-acting eQTLs for the global and local subnetworks (Fig. 4 whole blood; Fig. S7 muscle skeletal). Many of the CiTruss eQTLs for local subnetworks overlapped between the two tissue types (Figs. 5(c) and S10), suggesting both tissue-specific and shared genetic control of local subnetworks. Each *cis*-acting eQTL and the set of its eGenes and neighbors in the network overlapped well with an enhancer and the set of tissue-specific enhancer targets from EnhancerAtlas 2.0^20^, compared to the eGenes of the same eQTL found by Matrix eQTL (Fig. 5(d) whole blood; Fig. S8(c) muscle skeletal), providing evidence that the CiTruss gene networks and *cis*-acting eQTLs accurately capture the known gene modules controlled by *cis*-regulatory elements.

For the *trans*-acting eQTLs that CiTruss selected from the tissue-specific Type II blocks, we examined if they harbor transcription factors (TFs) for tissue-specific gene regulation and if their eGenes overlap with the known targets of TFs in hTFtarget ^21^ for whole blood and in JASPAR ^22,23^ for muscle skeletal. While in whole blood, the 23 *trans*-acting eQTLs and their eGenes did not have matching TF-target pairs, in muscle skeletal, three out of 21 *trans*-acting eQTLs had a TF nearby, all of their eGenes were targets of the given TF, and all of the three TFs and their eGenes were known skeletal-muscle genes (Table 3). Thus, these *trans*-acting eQTLs with direct effects on the eGenes were well supported as harboring a tissue-specific regulator by the known biology of muscle skeletal tissue.

### LG×SM AIL mouse gene networks and genetic diversity

We applied CiTruss combined with Matrix eQTL to the data from three brain tissue types, HIP, PFC, and STR, of the LG×SM AIL mice ^14^. The mouse and human gene networks shared similarities in the overall structure, but had several major differences (Figs. 6(a)-(b) PFC; Figs. S13(a)-(b) STR; Figs. S14(a)-(b) HIP). The mouse gene networks had substantially fewer connected genes (31.8% PFC; 30.8% STR; 31.9% HIP; Fig. S11), compared to the human gene networks (76.6% whole blood; 68.0% muscle skeletal), which led to fewer local subnetworks. The mouse gene network had a set of many global subnetworks that were extensively connected to local subnetworks, unlike the human gene networks with a single global subnetwork separated from local subnetworks. These differences arise from the difference in genetic diversity between the two populations. With fewer genetic variants and with differences in fewer phenotypes in behavior that originated from the two founders, the mouse gene network is more compact with fewer connections and as we discuss below, is more focused on brain function that is likely to explain the behavior differences in mice.

**Figure 6.**
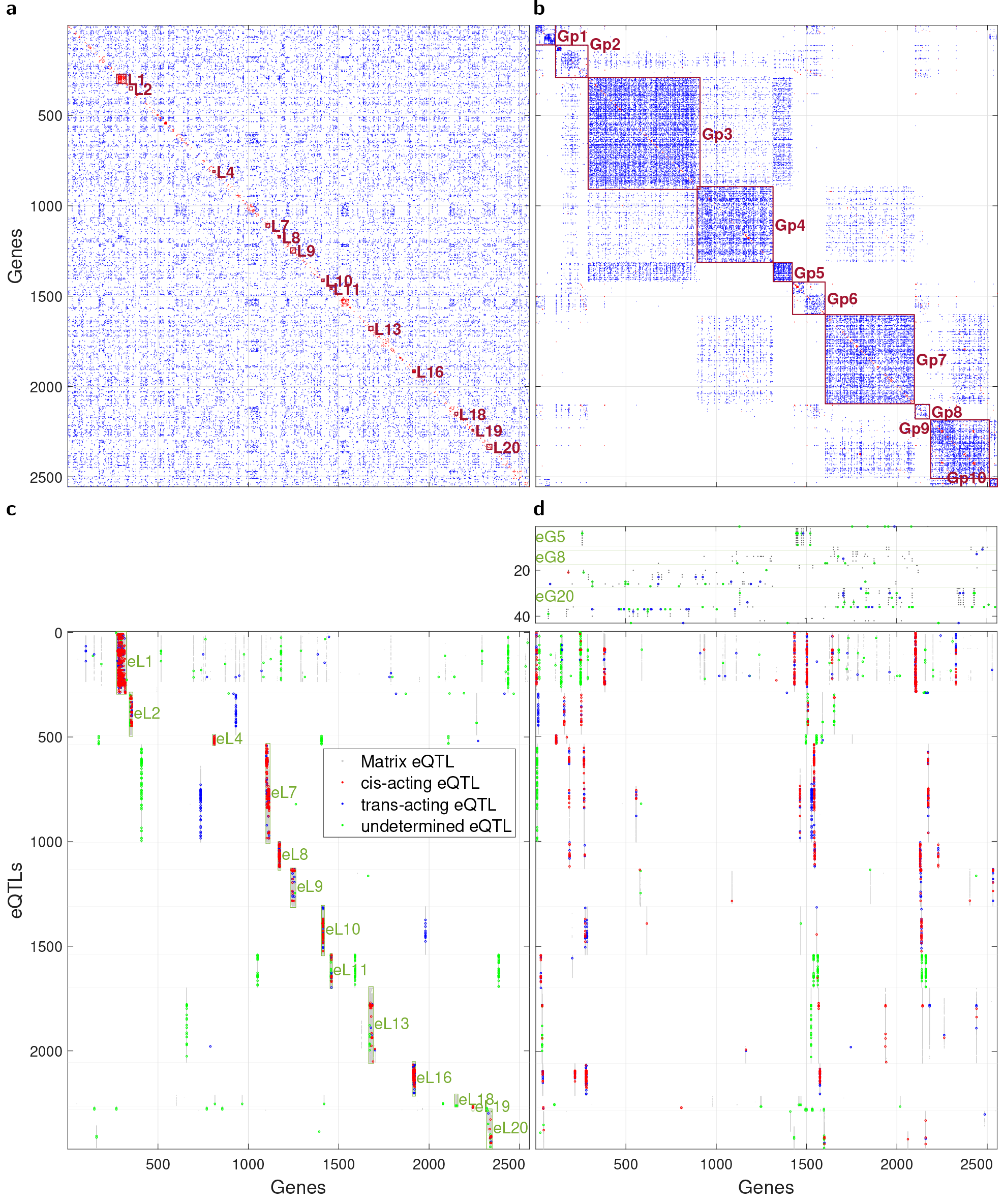
CiTruss gene network and eQTLs from LG×SM AIL mouse data for PFC tissue. (a) CiTruss gene network when genes are ordered with genome positions. Local subnetworks are shown as L1, etc. Isolated genes are not shown. (b) CiTruss gene network when genes are re-ordered after clustering. Global subnetworks are shown as Gp1, etc. (c) Hotspot eQTLs from Matrix eQTL (more than 5 eGenes) against the gene network in Panel (a). Type I blocks from hotspot regions (eL1, etc.) and local subnetworks (L1, etc.) are shown. (d) Hotspot eQTLs from Matrix eQTL against the gene network in Panel (b). Type II blocks (top) from hotspot eQTLs (eG1, etc.) and global subnetworks (Gp1, etc.), and Type I blocks (bottom). In Panels (c) and (d), *cis*-acting, *trans*-acting, and undetermined eQTLs from CiTruss are overlaid. See Tables S2 and S3 for locations of hotspot regions and subnetworks.

### LG×SM AIL mouse tissue-specific gene networks and pleiotropy

As with the human global subnetworks, nearly all mouse global subnetworks were enriched for genes annotated with tissue-specific brain GO terms (Table 2). However, unlike the two unrelated GTEx tissue types, the three related brain tissue types in mice shared a large number of genes in the sets of global subnetworks (1,613 between PFC and STR, 1,806 between PFC and HIP, and 1,738 between HIP and STR, out of 2,555 PFC, 2,593 STR, and 2,651 HIP), but these genes were organized into different global subnetworks in each tissue type. These observations from the GTEx human and AIL mouse gene networks suggest a hierarchical organization of global subnetworks across tissue types: at a higher level, unrelated tissue types do not share genes, whereas at a lower level, related tissue types share genes that are wired as different global subnetworks.

**Table 2:** GO terms enriched in CiTruss global subnetworks from LG×SM AIL mouse data.

| Subnet |  | Over | FDR | GO |  |
| --- | --- | --- | --- | --- | --- |
| ID | Size | lap |  | Term | Size |
| PFC |  |  |  |  |  |
| Gp1 | 109 | 10 | $9.1\times10^{-6}$ | BP Amino acid transport | 127 |
| Gp2 | 179 | 34 | $3.1\times10^{-2}$ | Nervous system development | 2068 |
| Gp3 | 619 | 70 | $4.1\times10^{-19}$ | Regulation of trans-synaptic signaling | 597 |
| Gp4 | 420 | 44 | $1.9\times10^{-10}$ | Regulation of trans-synaptic signaling | 597 |
| Gp5 | 105 | 4 | $9.1\times10^{-3}$ | Dendritic spine morphogenesis | 21 |
| Gp6 | 180 | 8 | $1.7\times10^{-3}$ | Axon ensheathment | 125 |
| Gp7 | 493 | 100 | $4.5\times10^{-10}$ | Nervous system development | 2068 |
| Gp8 | 80 | 5 | $4.4\times10^{-2}$ | Circadian regulation of gene expression | 74 |
| Gp9 | 325 | 32 | $4.3\times10^{-6}$ | Regulation of trans-synaptic signaling | 597 |
| Gp10 | 44 | 8 | $4.1\times10^{-10}$ | Translation at synapse | 48 |
| STR |  |  |  |  |  |
| Gs1 | 102 | 9 | $2.6\times10^{-2}$ | BP Regulation of synapse organization | 304 |
| Gs2 | 57 | 10 | $7.6\times10^{-8}$ | Synaptic vesicle cycle | 156 |
| Gs3 | 365 | 10 | $1.2\times10^{-3}$ | Regulation of neuronal synaptic plasticity | 74 |
| Gs4 | 93 | 7 | $2.2\times10^{-3}$ | Axon ensheathment | 125 |
| Gs5 | 658 | 52 | $1.1\times10^{-7}$ | Regulation of trans-synaptic signaling | 597 |
| Gs6 | 226 | 24 | $1.2\times10^{-7}$ | Synaptic signaling | 437 |
| Gs7 | 164 | 20 | $2.5\times10^{-8}$ | Anterograde trans-synaptic signaling | 365 |
| Gs8 | 173 | 48 | $9.5\times10^{-4}$ | Regulation of multicellular organismal process | 3264 |
| Gs9 | 40 | 9 | $1.4\times10^{-4}$ | Anterograde trans-synaptic signaling | 365 |
| Gs10 | 148 | 20 | $4.5\times10^{-4}$ | Protein modification by small protein conjugation/removal | 705 |
| Gs11 | 107 | 20 | $1.2\times10^{-7}$ | Regulation of trans-synaptic signaling | 597 |
| Gs12 | 347 | 27 | $2.1\times10^{-6}$ | Synaptic signaling | 437 |
| Gs13 | 94 | — | — | — | — |
| HIP |  |  |  |  |  |
| Gh1 | 97 | 7 | $1.1\times10^{-2}$ | BP Cytoplasmic translation | 113 |
| Gh2 | 68 | 8 | $1.3\times10^{-5}$ | Oxidative phosphorylation | 113 |
| Gh3 | 74 | — | — | — | — |
| Gh4 | 1027 | 95 | $1.0\times10^{-19}$ | Regulation of trans-synaptic signaling | 597 |
| Gh5 | 542 | 50 | $4.4\times10^{-15}$ | Synaptic signaling | 437 |
| Gh6 | 57 | — | — | — | — |
| Gh7 | 232 | 12 | $1.5\times10^{-4}$ | Synaptic vesicle cycle | 156 |
| Gh8 | 200 | 56 | $1.1\times10^{-2}$ | Transport | 3582 |
| Gh9 | 112 | 17 | $7.3\times10^{-4}$ | Monoatomic cation transport | 714 |
| Gh10 | 245 | 12 | $4.7\times10^{-6}$ | Axon ensheathment | 125 |

**Table 3:**
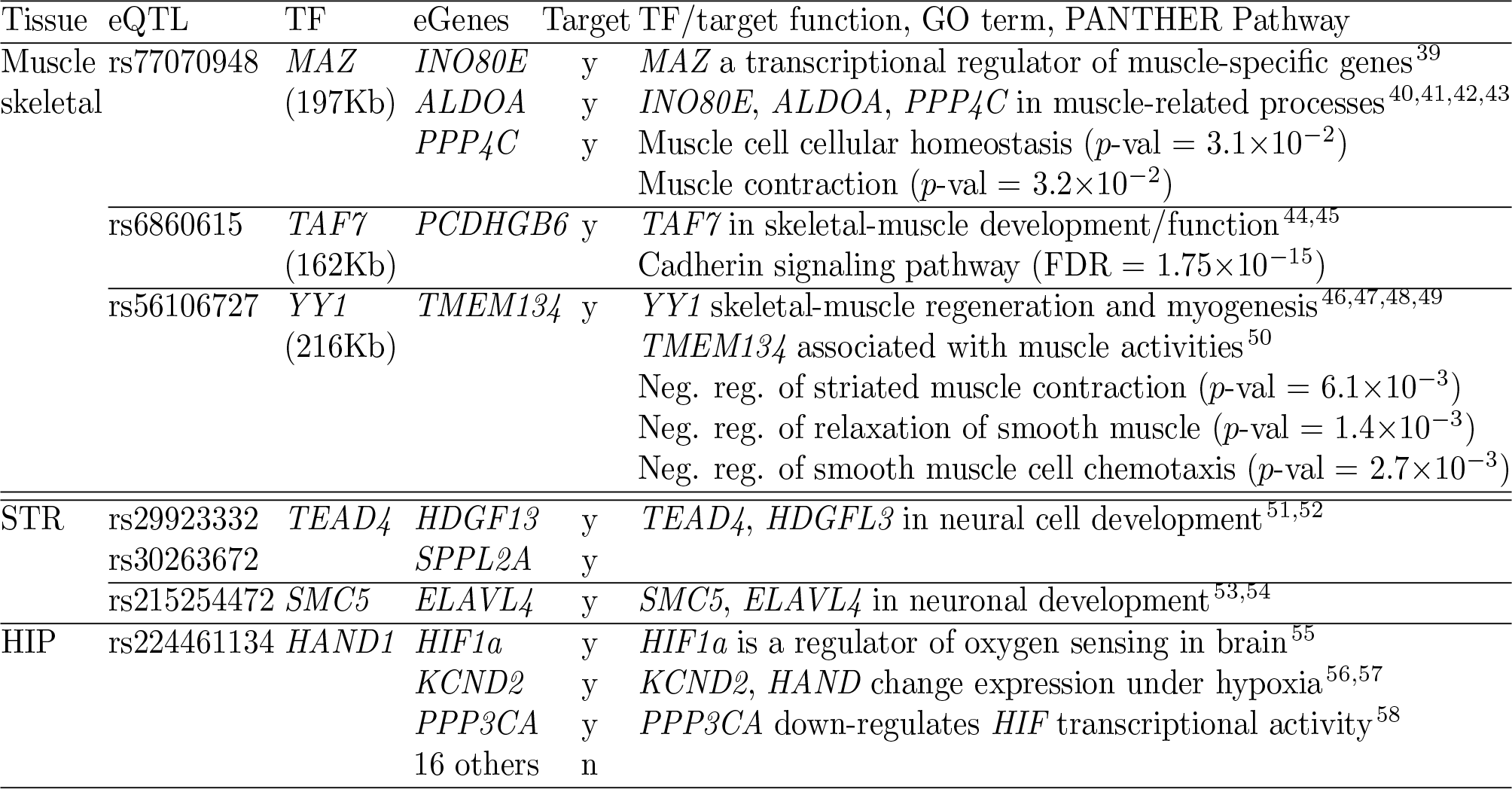
Overlap between *trans*-acting eQTL-eGenes and TF-targets.

Examining hotspot eQTLs with more than five eGenes from Matrix eQTL against CiTruss gene networks revealed that tissue-specific pleiotropy is organized around the local and global subnetworks: different tissue types often shared hotspot eQTLs for local subnetworks but not for global subnetworks. As with the GTEx eQTLs, the mouse hotspot eQTLs appeared as Type I or Type II blocks, but in a slightly different form (Figs. 6(c)-(d) PFC; Figs. S13(c)-(d) STR; Figs. S14(c)-(d) HIP). The three tissue types often had Type I blocks with the same hotspot regions and local subnetworks (Fig. S15(a), Table S2), but with different global subnetworks. Multiple Type II blocks were found in mice, each with 2 to 27 eQTLs in linkage disequilibrium (Table S3) and each affecting different global subnetworks, compared to a single block in humans. Long-range linkage disequilibrium was not observed in the mouse Type II blocks, suggesting that the mice did not undergo selection during the 56 generations of intercross.

### LG×SM AIL mouse eQTLs with direct effects

Nearly all of *cis*-acting eQTLs and the large portion of *trans*-acting eQTLs were found within 10Mb of their eGenes (Fig. S12), although distances to their eGenes tended to be larger in mice than in humans because of longer linked regions. Similar to the GTEx eQTLs, CiTruss disentangled Type I and II blocks to mostly *cis*-acting and *trans*-acting eQTLs for the local subnetworks, where many eQTLs overlapped across tissue types (Figs. S15(b)-(d)), and *trans*-acting and undetermined eQTLs for the global subnetworks (Fig. 6 PFC; Fig. S13 STR; Fig. S14 HIP). For the *trans*-acting eQTLs from Type II blocks, we looked for TFs in the genome neighborhood and the targets of the TF in the TFLink database ^24^ that overlap with the eGenes. We focused on Type II blocks, as these eQTLs were in relatively small linked regions. Across all tissue types, 15 out of 57 *trans*-acting eQTLs were located within 500Kb from a TF, five of these 15 eQTLs had eGenes overlapping with the TF targets, and four of these five eQTLs had a TF and targets that are known to be involved in brain function (Table 3).

## Discussion

We introduced CiTruss, a statistical framework for reconstructing gene networks perturbed by *cis*-acting and *trans*-acting eQTLs from allele-specific expression and SNP genotype data. Our analysis of the GTEx and AIL mouse data with CiTruss showed that gene networks and eQTLs should be identified in a single statistical analysis, since the genetic diversity in data determines the diversity in gene network modules that can be recovered from the data. The GTEx population had a large number of SNPs that perturbed the expression of many genes to cause variability in a broad range of phenotypes. CiTruss leveraged this high expression variability to reveal gene networks with a large number of local subnetworks. In contrast, the mouse gene networks from CiTruss had fewer connected genes and local subnetworks, but many global subnetworks with mostly brain-related genes that likely explain the behavioral differences between the two founder strains.

In addition, our results showed that identifying eQTLs along with gene networks allows us to gain new insights into various aspects of gene regulation controlled by eQTLs. CiTruss revealed that the pleiotropic effects of eQTLs, a hierarchy in tissue-specific gene regulation, and local and long-range linkage disequilibrium among eQTLs are structured around local and global subnetworks, and disentangled pleiotropic effects into *cis*-acting and *trans*-acting eQTLs for local subnetworks and *trans*-acting and undetermined eQTLs for global subnetworks. One future direction would be to determine a high-resolution map of hierarchical organization of the global subnetworks across tissue types, by applying CiTruss to eQTL data for a wide variety of tissue types from humans ^13^, farm animals ^16^, and rats ^25^. To further investigate the role of local and long-range linkage disequilibrium of eQTLs in gene regulation, we could compare these eQTLs with Hi-C data ^26^ to see if long-range chromatin interactions overlap with eQTLs in long-range linkage disequilibrium and if topologically associating domains overlap with the regions of linked eQTLs that contain local subnetworks.

The CiTruss framework could be extended in several ways. As population structure is a well-known confounding factor in eQTL mapping ^27,28^, to account for sample relatedness, we could extend the gene network parameter in the CiTruss model to the Cartesian product of two networks ^29^, one over genes and the other over samples. Another future direction is to combine SNP and expression data with trait data, to model a cascade of perturbation from genetic variants to gene expression to traits ^10^, by stacking a single-level conditional Gaussian graphical model for the probability distribution of traits given expression, on top of the multi-level model of CiTruss for the probability distribution of expression given genetic variants. Finally, CiTruss could be adopted to reconstruct gene networks from expression data obtained from single-cell RNA-seq after CRISPR genome editing ^30,31^, replacing SNP perturbation with CRISPR perturbation.

## Material and Methods

We describe the probability model, learning algorithm, and inference method of our statistical framework CiTruss, and provide the details of the experimental set-up.

### Multi-level conditional Gaussian graphical models

We introduce our multi-level conditional Gaussian graphical model for modeling a gene network perturbed by *cis*-acting and *trans*-acting eQTLs. We extend the existing single-level conditional Gaussian graphical model for learning a gene network perturbed by eQTLs to a multi-level model with a two-level hierarchical structure. In the top level, both alleles of a gene interact with both alleles of a neighboring gene in the network and are affected by both alleles of SNPs or *trans*-acting eQTLs. In the bottom level, the two alleles of each gene act independently in allele-specific manner and are affected by SNP allele on the same haplotype in *cis*.

Let 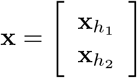, where 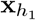, 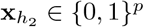 with two alleles 0 and 1 denote alleles at *p* SNP loci on two haplotypes *h*_1_ nd *h*_2_. Let 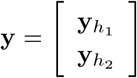, where 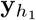, 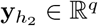 represent the expression levels of the two alleles of *q* genes from haplotypes *h*_1_ and *h*_2_. Then, the multi-level conditional Gaussian graphical model for a gene network perturbed by *cis*-acting and *trans*-acting eQTLs (Fig. 1(a)) is

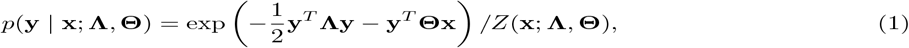

where the 2*q* × 2*q* positive definite matrix **Λ** is a parameter representing a network over 2*q* gene alleles, **Θ** ∈ ℝ ^2*q*×2*p*^ is a parameter representing the perturbation of this network by eQTLs, and 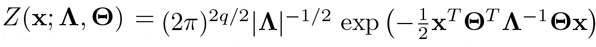 is a normalization factor, known as a partition function, that ensures the probability density function integrates to 1. Both **Λ**and **Θ** have a multi-level structure defined as the sum of two component matrices:

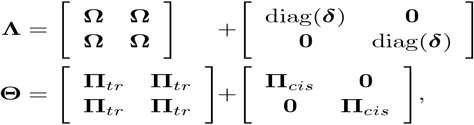

where **Ω** is a *q* × *q* positive-semidefinite matrix with zeros in the diagonal representing a gene network over *q* genes, diag(***δ***) is a diagonal matrix with *q* × 1 vector ***δ*** of positive values that models the variability in the expression of two alleles, and **Π**_*tr*_ and **Π**_*cis*_ ∈ ℝ^*q*×*p*^ represent *trans*-acting and *cis*-acting eQTL effects, respectively. A non-zero value in the (*i, j*)th element of **Ω**represents an edge between gene *i* and gene *j*, and a non-zero value in the (*i, j*)th element of **Π**_*cis*_ and **Π**_*tr*_ represents SNP *j* affecting the expression of gene *i* in *cis* and *trans*, respectively.

The first component matrices of **Λ**and **Θ** correspond to the top-level model, where the expression of both alleles of each gene is influenced by the expression of both alleles of neighboring genes in the network **Ω**and by *trans*-acting eQTLs in **Π**_*tr*_. The four blocks of **Ω**force the expression of both alleles of each gene to be the same, and the four blocks of **Π**_*tr*_ model *trans*-acting eQTL effects of 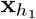 and 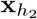 on both 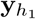 and 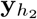. The second component matrices of **Λ**and **Θ** correspond to the bottom-level model, where the expression of the two alleles of each gene varies independently in an allele-specific manner and is affected by *cis*-acting eQTLs. The two diagonal blocks of diag(***δ***) model the allele-specific expression variability and the two diagonal blocks of **Π**_*cis*_ model *cis*-acting eQTL effects of 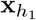 on 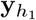, and 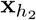 on 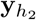.

### Hybrid model for partially observed allele-specific expression

If the coding region of a gene has no heterozygous loci, the two alleles of the gene are indistinguishable, and only the total expression of both alleles can be measured. When allele-specific expression is measured only for a subset of genes, the multi-level model in Eq. (1) collapses to a hybrid model that consists of a single-level model for genes with only total expression and a multi-level model for genes with observed allele-specific expression. The multi-level model collapses differently for different individuals, since each individual has a unique genome sequence with its own set of heterozygous loci. The following theorem states that this hybrid model for a given individual is also a conditional Gaussian graphical model (illustration in Figs. 1(b)-(c); proof in Appendix).

#### Theorem 1.

**(Hybrid model)** *For a given individual, let H* ⊂ {1, …, *q*} *and S* = {1, …, *q*} − *H be the subsets of genes with observed and unobserved allele-specific expression, respectively. Let* **y**^*H*^ *denote a subvector of* **y** *for genes in H and* 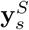 *a subvector of total expression* 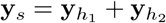 *for genes in S. Let* 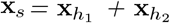 *represent genotypes. Then, the multi-level model in Eq. (1) for this individual collapses to a hybrid of multi-level model for* **y**^*H*^ *and single-level model for* **y**^*S*^. *The resulting hybrid model is a conditional Gaussian graphical model for* 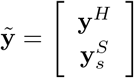, *given* 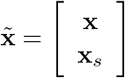:

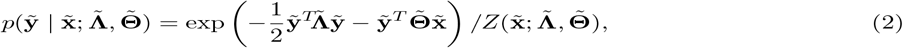

*where*

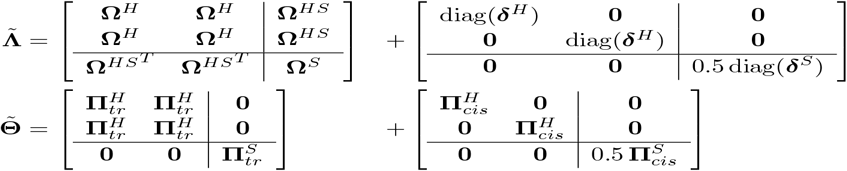

*are the network and perturbation parameters. In* 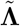, **Ω**^*H*^, **Ω**^*S*^, *and* **Ω**^*HS*^ *are the submatrices of* **Ω***that consist of the* (*i, j*)*th element of* **Ω***for all i, j* ∈ *S, i, j* ∈ *H, and i* ∈ *H, j* ∈ *S, respectively. In* 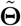, 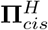 *and* 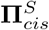 *are the submatrices of* **Π**_*cis*_ *that consist of the ith row of* **Π**_*cis*_ *for all i* ∈ *H and i* ∈ *S, respectively*. diag(***δ***^*H*^), diag(***δ***^*S*^), 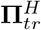, *and* 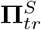 *are defined similarly*.

In the hybrid model above, the network and eQTLs for the multi-level model are in the upper-left blocks of 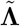 and 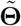, and those for the single-level model are in the lower-right blocks of 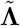 and 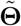. For each gene in *S* with only total expression, the two nodes in the network **Λ**for its two alleles collapse into a single node in 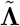, and the edges between the given gene and its neighbors in the network also collapse (collapse from Fig. 1(b) to Fig. 1(e) for one individual, and from Fig. 1(c) to Fig. 1(f) for another individual). The *cis*-acting and *trans*-acting eQTLs in **Θ** that affect the genes in *S* collapse to eQTLs in 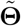. In the extreme case where allele-specific expression is unobserved for all genes, the multi-level model collapses to a single-level model for **y**_*s*_ given **x**_*s*_:

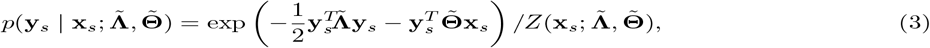

where 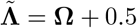 diag(***δ***) and 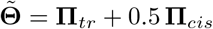. Even when allele-specific expression levels are available, only heterozygous variants contain information to determine whether a variant is affecting gene expression in *cis* or *trans*. If all variants are homozygous, even when allele-specific expression is measured for all genes, the multi-level model for the given individual collapses to Eq. (3).

### Sum-difference model for total and differential expression of two alleles

Estimating the multi-level model parameters is prohibitively expensive, since the model has a large network parameter and collapses differently for each individual. Here, for efficient parameter estimation, we apply a linear transformation to the random variables of the multi-level model to obtain a sum-difference model that factorizes into two single-level models: a sum model for the total expression perturbed by eQTLs and a difference model for the differential expression of two alleles controlled by *cis*-acting eQTLs. The following theorem shows the sum-difference model and its factorization (illustration in Fig. 1(d); proof in Appendix).

#### Theorem 2.

**(From multi-level model to sum-difference model)** *In addition to sum variables* **y**_*s*_ *and* **x**_*s*_, *we define diffe rence va riab les* 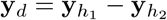 *and* 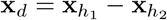. *We apply the transformation of random variables from* 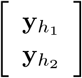 *to* 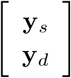 *the multi-level model in Eq. (1)*. *Then, the resulting probability distribution is a sum-difference model that factorizes as follows:*

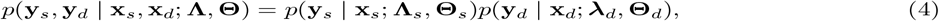

*where the first probability factor is a sum model for sum variables*

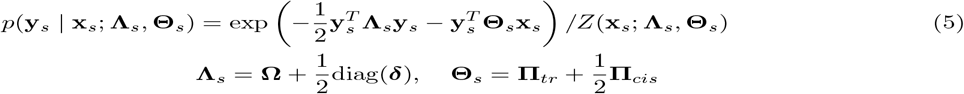

*and the second probability factor is a difference model for difference variables*

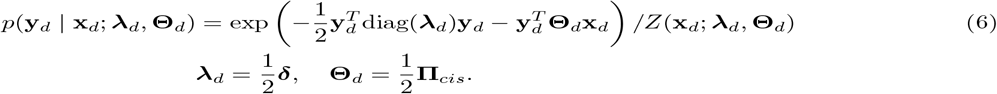

*The difference model in Eq. (6) further factorizes into q single-gene difference models for q genes as follows:*

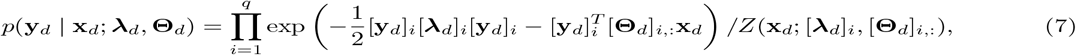

*where* [**a**]_*i*_ *represents the ith element of vector* **a** *and* [**C**]_*i*,:_ *represents the ith row of matrix* **C**.

In the sum-difference model above, only the difference model contains information on allele-specific expression, and the sum model is defined entirely in terms of the total expression of two alleles. It is straightforward to show that the sum-difference model achieves maximum likelihood at the same parameter value as the multi-level model. The following corollary of Theorem 2 shows that the same transformation can be used to obtain a sum-difference model and its factorization from the hybrid model (illustration in Figs. 1(e)-(f); proof in Appendix).

#### Corollary 1.

**(From hybrid model to sum-difference model)** *Applying the same transformation of random variables as in Theorem 2 to the hybrid model in Eq. (2), we obtain a sum-difference model that factorizes as*

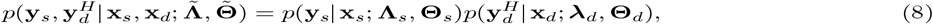

*where the sum and difference models are given as the following single-level models:*

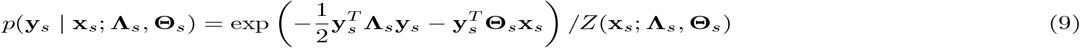

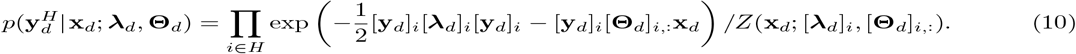

The corollary above shows that in sum-difference model, only the difference model changes with unobserved allele-specific expression. In other words, the sum models derived from the multi-level and hybrid models are identical. The difference model from the hybrid model is a reduced form of the difference model from the multi-level model in that it contains the single-gene probability factors only for genes with allele-specific expression.

### Parameter estimation using sum-difference model

We estimate the model parameters based on sum-difference model (see Appendix for detail). With its factorization, we estimate each of the sum and difference model parameters efficiently for a large number of SNPs and genes, using Mega-CGGM ^32,10^, the existing efficient optimization method for single-level model. Parameter estimation is significantly more efficient with the sum-difference model than with the multi-level model, owing to the local changes to the difference model with unobserved allele-specific expression. In the sum-difference model, the evaluation of the partition function, which involves computing the matrix inverse and determinant with cost *O*(*q*^3^), needs to be performed only once to be reused for all individuals in each iteration of the optimization. In the multi-level model, this computation needs to be performed separately for each individual in each iteration of optimization for a larger matrix of up to size 2*q* × 2*q*. This reduction in per-iteration time cost from *O*(*n*(2*q*)^3^) to *O*(*q*^3^) leads to orders-of-magnitude speed-up empirically.

### Inferring downstream effects of eQTLs in gene network

While the perturbation parameter **Θ** represents eQTLs with direct effects on gene expression, such direct perturbation effects can propagate through the gene network **Λ**to affect the expression of other genes indirectly. To infer such indirect eQTL effects, we re-write Eq. (1) as a Gaussian distribution

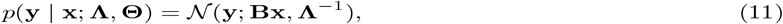

where the 2*q* × 2*p* matrix

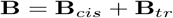

with

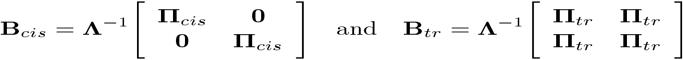

represents the aggregate effects of SNPs on gene expression that include both direct and indirect effects. A non-zero value in the (*i, j*)th element of **B** but a zero in the (*i, j*)th element of **Θ** indicates the *i*th gene is not perturbed directly by the *j*th SNP but is indirectly perturbed due to the downstream effects of the *j*th SNP directly perturbing other genes in the network. The overall indirect effects **B** can be decomposed into two parts **B**_*cis*_ and **B**_*tr*_, each induced by *cis*-acting eQTLs and by *trans*-acting eQTLs. If the gene network has no edges and is a diagonal matrix **Λ**= λ**I**_2*q*_, then Eq. (11) reduces to a scaled multivariate regression model.

### Re-analysis of eQTLs from the existing methods with CiTruss

CiTruss can be combined with the existing methods such as Matrix eQTL, TReCASE, WASP, and RASQUAL that identify SNP-gene pairs with statistically significant association. The pleiotropic effects of an eQTL on many genes with correlated expression found by these existing methods could indicate few eQTLs that perturb few genes directly and affect downstream genes in the pathway indirectly. Given such pleiotropic effects, our approach can be used to identify a network over these genes and eQTLs with direct effects on genes in the network. In such re-analysis, during parameter estimation, the sum model is constrained so that eQTLs are a subset of the eQTLs found by the existing methods and then the difference model is constrained such that *cis*-acting eQTLs are a subset of the eQTLs from the sum model. This is implemented by modifying Mega-CGGM such that an active set is always a subset of the eQTLs from the existing method.

### Generating simulated data

To evaluate the various methods on datasets with known ground truth, we simulated data from known eQTLs and gene networks. We generated SNP genotype data at each of *p* = 100 loci for *n* = 200 individuals by sampling genotypes from a multinoulli distribution over the number of minor alleles {0, 1, 2} with probabilities 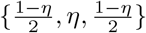, where *η* is the frequency of heterozygous genotypes in the population at a given locus. We simulated three sets of genotype data, each with *η* = 0.05, 0.25, and 0.45, and set the alleles on the haplotypes to be consistent with these genotypes. We then simulated allele-specific expression data from our model with known ground-truth gene network and *cis*-acting/*trans*-acting eQTLs set as follows. We generated a gene network **Ω**over *q* = 40 genes, assuming four gene modules each with 10 genes and assuming 40% non-zero elements within each module and 2% non-zero elements between modules. The values of the non-zero elements were randomly generated from Uniform[−4, 4] for edges within modules and Uniform[−1, 1] for edges between modules. The diagonal elements of **Ω**were set such that the minimum eigenvalue is 0.1 and the maximum diagonal element is not 70% larger than the maximum off-diagonal element. For **Π**_*cis*_ and **Π**_*tr*_, we selected 5% of all pairs of genes and SNPs to have non-zero elements in either **Π**_*cis*_ or **Π**_*tr*_ with equal probability and set their values to random samples from Uniform[−8, 8].

Given the SNP data and ground-truth parameters, we generated allele-specific expression data from our multi-level model. To mimic partially observed allele-specific expression, we kept the allele-specific expression for a given gene in the population at frequency *ζ*(*ζ*= 1.0, 0.9, 0.7, and 0.5 in each dataset) and replaced the allele-specific expression levels for the rest of the individuals with the total expression levels of the two alleles. For each combination of *η* and *ζ*, we generated five datasets and report results averaged over these datasets. For methods such as TReCASE, RASQUAL, and WASP that require allele-specific read counts, we transformed the simulated data above to count data through inverse log normalization, while ensuring the minimum and maximum of the read counts for each gene across individuals match those of log_2_ of transcripts per million (TPM) in the GTEx whole blood samples.

### Comparison of methods on simulated data

In simulated data analysis, for TReCASE and Matrix eQTL, as the authors of TReCASE suggested, we included seven covariates for each sample: log of total expression in TPM, top three principal components of expression data in TPMs, and top three principal components of expression data in log of TPM. We excluded covariates with variance less than 10^−4^ across samples. For WASP, we ran the combined haplotype test, using the overdispersion parameters estimated from data for both association and allele-specific tests.

In re-analysis of the eQTLs identified by the existing methods with CiTruss, the sum model was constrained to select eQTLs from *cis*-acting and *trans*-acting eQTLs identified by TReCASE, from *cis*-acting eQTLs identified by RASQUAL or WASP, and from eQTLs identified by Matrix eQTL. Then, the difference model was constrained to select *cis*-acting eQTLs from the eQTLs selected by the sum model. The ground-truth eQTLs with indirect effects were obtained from Eq. (11) with the ground-truth parameters.

### Preprocessing GTEx data

We downloaded phased genotype data for 46,526,292 SNPs and allele-specific expression data for 56,200 genes from the whole blood of 670 individuals and from the muscle skeletal of 706 samples from the GTEx v8 repository ^13^. We analyzed 1,175,808 SNPs for whole blood and 1,172,754 SNPs for muscle skeletal, after excluding indels and SNPs with missing genotype calls and with MAF< 0.05, and removing SNPs with *r*^2^ > 0.8 using PLINK 1.9^33^ with a 1000kb sliding window. For the total expression of both alleles, we included in our analysis 10,636 genes for whole blood and 12,475 genes for muscle skeletal with sample variance of the total expression of two alleles in TPM > 100. For allele-specific expression, we included 9,240 genes for whole blood and 10,513 genes for muscle skeletal whose sample variance of the differences in the expression of two alleles is at least 10 and whose frequency of observed allele-specific expression is at least 0.02. In the fully processed data, the minimum frequency of heterozygous genotypes was 0.024 for whole blood and 0.026 for muscle skeletal.

### Preprocessing LG×SM AIL mouse data

We prepared phased genotype and allele-specific expression data from the raw genotyping-by-sequencing data and RNA-seq reads from HIP, PFC, and STR tissues of the LG×SM AIL mice ^14^. We called and phased genotypes for 1,084 mice as follows. We removed SNPs with MAF*<*0.005 and with reads in fewer than 20% of the samples using bcftools ^34^. Then, we called genotypes from the genotype likelihoods and filled in missing genotypes using Beagle 4.1^35^ (window size 300bp, step size 50bp for sliding windows, tolerance 0.04 for convergence, and the maximum number of iterations 10) with the mouse genetic map. We phased and imputed the resulting 50,568 SNPs with MAF> 0.1, using Beagle 4.1^36^ (window size 70kb, step size 32kb for sliding window, tolerance 0.03 for convergence, and the maximum number of iterations 10) with the LG and SM founder genomes as a reference panel. We excluded SNPs deviating from Hardy-Weinberg equilibrium (*p*-value ≤ 7.62 × 10^−6^).

We performed allele-specific expression quantification from RNA-seq reads as follows. For each sample, we aligned reads to the personalized diploid transcriptome constructed from the phased genotypes and mouse transcript sequences (M25 GRCm38.p6) using Bowtie. Bowtie was run with options to report only the alignments in the best stratum with at most three mismatches and to suppress reads with more than 100 valid alignments. We then quantified allele-specific expression using EMASE-Zero Model 2 (tolerance 10^−8^ for convergence). Following the preprocessing steps of the original mouse study, we removed 7 samples from HIP, 5 from PFC, and 23 from STR with more than 1.5 standard deviation from the mean number of reads or the mean alignment rates, dropped 21 samples as contaminated outliers, based on clustering of samples with the top two principal components of total expression of two alleles from all tissue types, and corrected the labels of 18 HIP samples that were mislabeled as PFC or STR.

Then, for each tissue type, we selected samples that have both phased genotype and allele-specific expression data. For 239 such samples for HIP, 208 for PFC, and 189 for STR, we further processed the data as follows. After removing SNPs with *r*^2^ > 0.9999 in 3,000kb windows using PLINK 1.9^33^ and SNPs with MAF< 0.05, we obtained genotypes for 30,742 SNPs for HIP, 30,066 SNPs for PFC, and 29,139 SNPs for STR. For total expression of both alleles, we retained genes whose sample variance in the total expression measured as TPM is at least 5.0 in each tissue type, which led to total expression data for 8,298 genes for HIP, 8,030 genes for PFC, and 8,423 genes for STR. For allele-specific expression, we retained genes whose sample variance in the differences in the expression of two alleles is at least 5.0 and whose frequency of observed allele-specific expression in the population is at least 0.1. This led to allele-specific expression for 3,604 genes for HIP, 3,503 genes for PFC, and 3,571 genes for STR for a subset of the samples, and for the rest of the genes, allele-specific expression was assumed unobserved. In the fully processed data, the minimum frequency of heterozygous genotypes was 0.0084 for HIP, 0.0096 for PFC, and 0.0053 for STR, and all SNPs, except for two SNPs in each tissue type, had the frequency of heterozygous genotypes > 0.02.

### Comparison of methods on mouse and GTEx data

In our analysis of the GTEx data with Matrix eQTL, we included the seven covariates from the GTEx v8 repository: top five PEER factors ^13^ and top two principal components from genotype data. For the mouse data, we included seven covariates: log of total read counts, top three principal components of read counts, and top three principal components of log of read counts. We excluded covariates with variance < 10^−4^.

The total expression was used to estimate the sum model and the allele-specific expression was used to estimate the difference model. Genes that did not meet the criteria of the minimum frequency of observed allele-specific expression and the minimum sample variance were included in the sum-model estimation but not in the difference-model estimation. The eQTLs for these genes were called undetermined eQTLs, since it is not possible to determine whether they are *cis*-acting or *trans*-acting. In re-analysis of the eQTLs from Matrix eQTL with our approach, we constrained the sum model to identify eQTLs as a subset of the eQTLs from Matrix eQTL, and constrained the difference model to identify *cis*-acting eQTLs as a subset of the eQTLs selected in the sum model. We selected the regularization hyperparameters using BIC.

### Comparison of computation time

We implemented CiTruss as a C++ wrapper for Mega-sCGGM ^32^. We used the R implementation of TReCASE ^12^ and Matrix eQTL ^11^, the C implementation of RASQUAL ^6^, and the Python implementation of WASP combined haplotype test ^7^ provided by the authors. The experiments on the GTEx data were run on 128 cores of two AMD EPYC 7742 CPUs. The experiments on the simulated and mouse data were run on 8 cores of Intel(R) Xeon(R) Gold 6230.

### Comparison of *cis*-acting eQTLs with enhancer database

We downloaded the tissue-specific enhancer-gene interaction database from EnhancerAtlas 2.0^20^. We used the enhancer-gene interactions in human GM12878 lymphoblastoid cell line and skeletal muscle tissues. These interactions are computational predictions made by EAGLE ^37^ based on the information such as genomic locations and correlation between enhancer activity and gene expression levels ^20^. All interactions were between an enhancer and its target gene that are at most 1Mb apart. The database contained 78,796 enhancer-gene interactions (7,364 enhancers and up to 72 target genes per enhancer) for GM12878 lymphoblastoid cell line, and 39,708 enhancer-gene interactions (4,334 enhancers and up to 47 genes per enhancer) for human skeletal muscle. In each GTEx tissue type, we mapped each *cis*-acting eQTL to enhancers located within 100Kb, and the set of eGenes to the set of enhancer targets (hypergeometric test *p*-value < 0.05).

### Comparison of *trans*-acting eQTLs and eGenes with TFs and targets

We compared the pairs of *trans*-acting eQTLs and their eGenes with the pairs of TFs near the eQTLs and their targets from TF-target databases. We used hTFtarget ^21^ with tissue-specific TF-target relationship for the GTEx whole blood, and JASPAR database ^22^ curated by Harmonizome ^23^ for the GTEx skeletal muscle. For the AIL mice, we used TFLink database ^24^.

### Identifying local and global subnetworks in gene networks

To identify local and global subnetworks in gene networks estimated by CiTruss from the GTEx data, we applied hierarchical clustering with complete-linkage to the estimated gene networks. During clustering, we excluded the edges for gene pairs with less than 50 genes apart on the genome, to ensure the orderings of the genes in the local subnetworks are preserved. For the mouse gene networks, we applied hierarchical clustering with complete linkage to the sample correlation of log gene expression data, excluding the correlation between genes within 10Mb. The ordering of the genes given by hierarchical clustering was used to find the local and global subnetworks.

## Appendix

### Proofs of Theorems

We provide the proofs of the theorems and corollaries presented in the Material and Methods section.

#### Proof of Theorem 1

*Proof*. We define a (2|*H*| + |*S*|) × 2*q* matrix 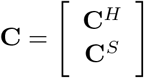, with the 2|*H*| × 2*q* matrix **C**^*H*^ and the |*S*| × 2*q* matrix **C**^*S*^. We define **C**^*H*^ as a submatrix of **I**_2*q*×2*q*_, taking rows corresponding to the alleles of genes in *H*, such that **y**^*H*^ = **C**^*H*^ **y**. We define **C**^*S*^ as a submatrix of [**I**_*q*×*q*_ **I**_*q*×*q*_], taking rows corresponding to genes in *S*, such that 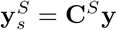. Let 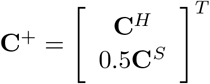 denote the Moore-Penrose inverse of **C**.

We re-write Eq. (1) in the form of Gaussian distribution and from this, we obtain the distribution of 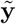, which is also Gaussian

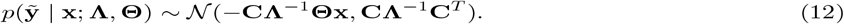

To prove the theorem, it is sufficient to show that Eq. (12) can be re-written as

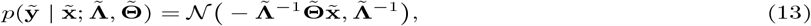

since expanding the quadratic term in Eq. (13) leads to the model in Eq. (2). Below, from Eq. (12), we derive the forms of 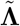, 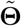, and 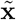 in Eq. (13). To obtain 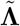 given in the theorem, we re-write the covariance matrix in Eq. (12) as follows:

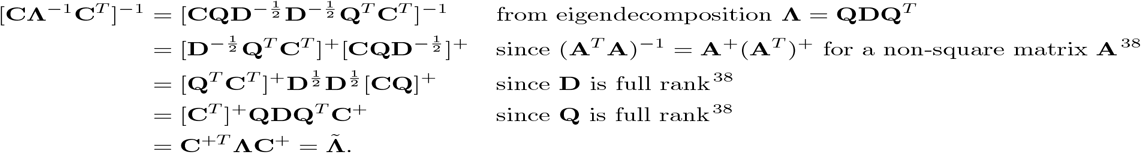

To obtain 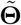 and 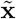 given in the theorem, we re-write the mean in Eq. (12) as follows

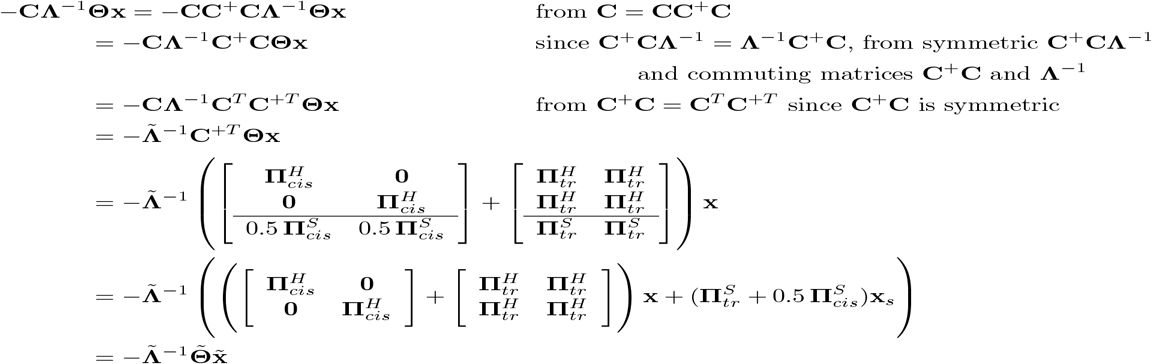

Thus, we have Eq. (13) with 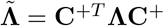, 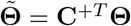, and 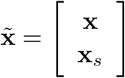.

The following corollary of Theorem 1 is a property of the hybrid model.

#### Corollary 2.

*The likelihood of the partially observed allele-specific expression data* 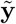 *given the collapsed model in Eq. (2) can be written in terms of the imputed data* 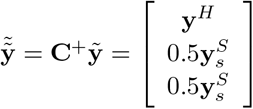, *where the expression levels* 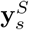 *are split evenly between the two alleles in the imputed data*.

*Proof*. We re-write the model in Eq. (2) as

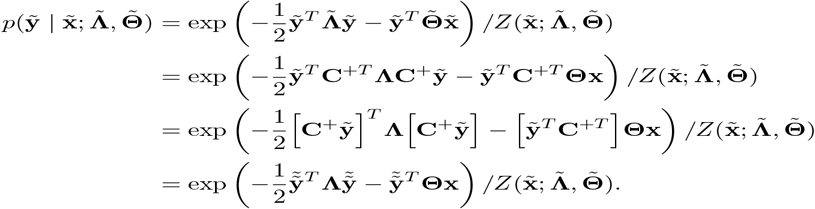

Thus, the likelihood 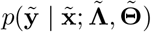 can be written in terms of the imputed data 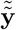.

#### Proof of Theorem 2

We prove the following lemma and use this lemma to prove Theorem 2.

#### Lemma 1.

*Let* 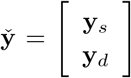*and* 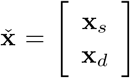. *Applying the change of random variables* 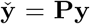, *where* 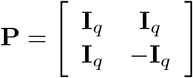, *to the multi-level model in Eq. (1), we obtain*

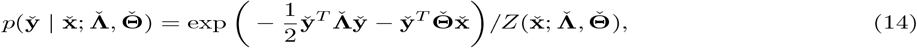

*where*

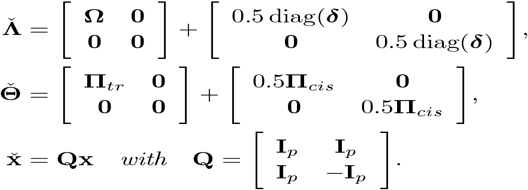

*Proof*. We apply the standard procedure for change of random variables as follows. Since 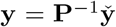 and 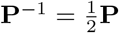, we substitute **y** with 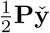 in Eq. (1) and multiply this with the Jacobian determinant of the inverse of the transformation 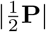:

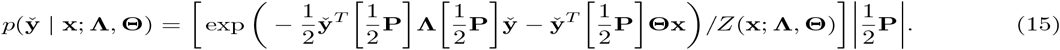

We visit each term in Eq. (15) to derive Eq. (14). For the first term in Eq. (15), it is straightforward to verify 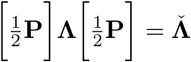. For the second term, we have

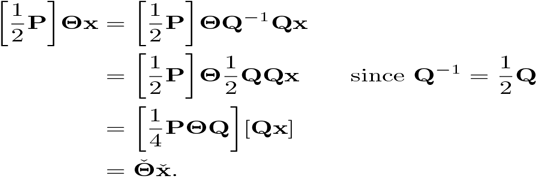

For the remaining terms in Eq. (15), we have

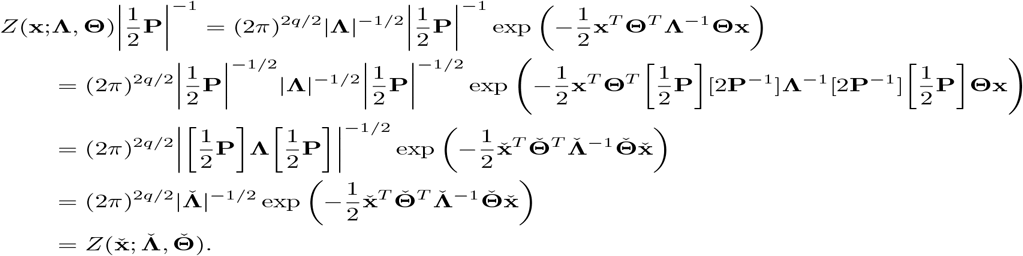

Combining the results above, we obtain Eq. (14).

The block-diagonal structure of the network matrix 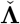 in Eq. (14) indicates that after the change of random variables, **y**_*s*_ and **y**_*d*_ become conditionally independent given 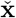. Below, we use this conditional independence to prove Theorem 2.

*Proof*. To obtain Eq. (4), we factor Eq. (14) in Lemma 1 into two probability models, each for **y**_*s*_ and **y**_*d*_. We first factor the numerator of Eq. (14) as follows:

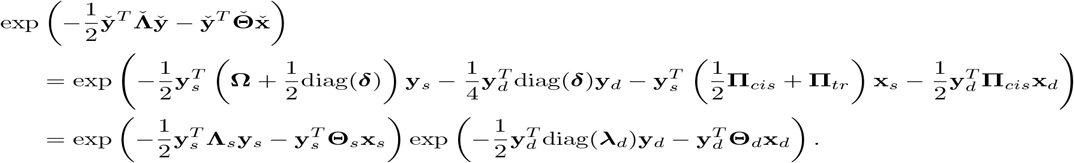

We then factor the denominator in Eq. (14) as

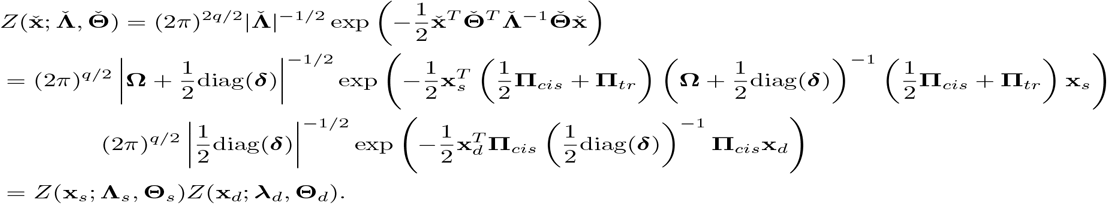

Combining the results on the numerator and denominator above, we have the decomposition in Eq. (4).

The difference model is further factorized into *q* single-gene models due to its diagonal inverse covariance matrix diag(**λ**_*d*_):

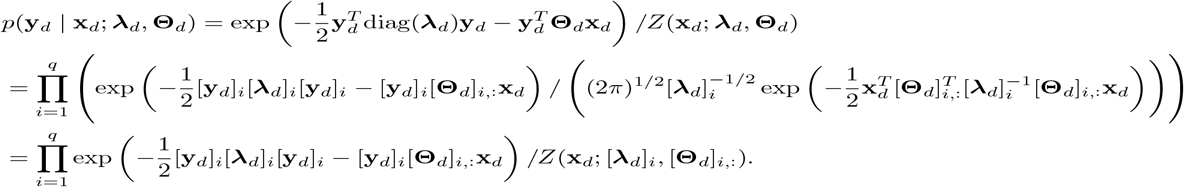

Notice that the transformation in Theorem 2 is an orthogonal transformation that involves rotation and reflection of the original random variable **y**.

**Proof of Corollary 1**

*Proof*. Applying the transformation 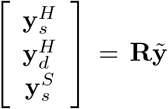, where 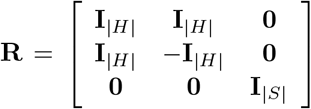, to the random variable 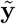of the hybrid model and reordering the random variables in 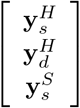 to 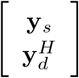 lead to the decomposition in Eq. (8).

### Parameter estimation with sum-difference model

We fit sum and difference models using sum and difference data derived from the original phased SNP data and allele-specific or total expression data. Given SNP allele data 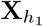, 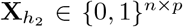 and allele-specific expression data 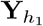, 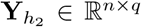 R for *n* individuals, we derive sum data, 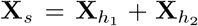and 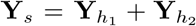, and difference data, 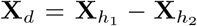and 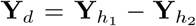. The (*i, j*)th element of **Y**_*d*_ for gene *j* and individual *i* is available, only if the (*i, j*)th element of 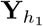 and 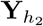 is available. We estimate the sum-difference model parameters **Λ**_*s*_, **Θ**_*s*_, **λ**_*d*_, and **Θ**_*d*_ by minimizing the *L*_1_-regularized negative log-likelihood of the sum and difference data:

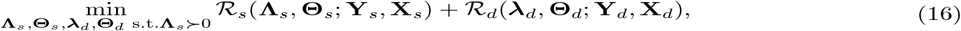

where

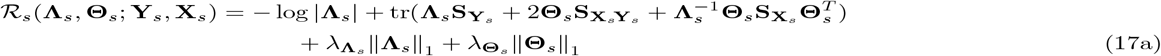

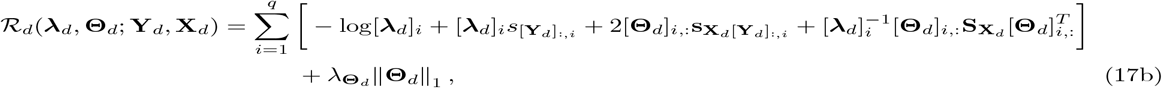

with covariance matrices 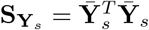, 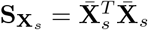, 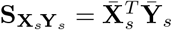, 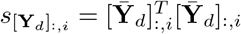, 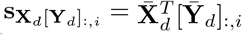, and 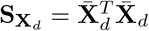 with mean-centered data 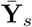, 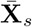, 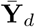, and 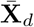. The *L*_1_ penalty || · || encourages the estimates to be sparse with many zeros in the parameter matrices, and has the property of selecting few SNPs with the most relevance to gene expression among many correlated SNPs in linked regions of genome. The regularization hyperparameters 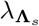, 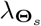, and 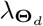 control the level of sparsity in the estimated parameters. We select the regularization hyperparameters with the Bayesian information criterion (BIC).

We solve Eq. (16) by sequentially minimizing Eq. (17a) and Eq. (17b) with Mega-CGGM. This amounts to first finding a gene network and eQTLs perturbing this network in the sum model and then determining whether these eQTLs are acting in *cis* in the difference model. When estimating the difference-model parameters, we constrain the set of non-zero elements in **Θ**_*d*_ to be a subset of the non-zero elements in the estimated **Θ**_*s*_ of the sum model, by constraining the active set to be this subset in each iteration in Mega-CGGM. This ensures that *cis*-acting eQTLs in the difference model are also eQTLs in the sum model. From the estimated **Λ**_*s*_, **Θ**_*s*_, **λ**_*d*_, and **Θ**_*d*_ and from Eqs. (5) and (6), we recover the multi-level model parameters as **Ω**= **Λ**_*s*_ − diag(**λ**_*d*_), ***δ*** = 2**λ**_*d*_, **Π**_*cis*_ = 2**Θ**_*d*_, and **Π**_*tr*_ = **Θ**_*s*_ − **Θ**_*d*_. Then, we determine *cis*-acting eQTLs based on the non-zero elements of **Π**_*cis*_ and *trans*-acting eQTLs based on the non-zero elements of **Π**_*tr*_ that are not *cis*-acting eQTLs. While the multi-level and sum-difference models give identical maximum likelihood estimates, the *L*_1_ regularization in Eq. (17a) and Eq. (17b) introduce bias that shrinks the estimates towards zero to induce sparsity. To account for this shrinkage effect, the non-zero elements of **Θ**_*s*_ that are *cis*-acting eQTLs are ruled out as *trans*-acting eQTLs, even though they may have both *cis* and *trans* effects.

## Acknowledgments

We thank Calvin McCarter and Gi Bum Kim for their contribution to the early stage of this work, Abraham Palmer for providing the RNA-seq reads for the AIL mice, Kathryn Roeder for her suggestion to pair up CiTruss with another statistical method, and Quasar Padiath for discussion on linkage disequilibrium. This work was supported by NIH-1R21HG010948, NSF-DBI2154089, and Sloan Research Fellowship.

## Data and Code Availability

Contact the authors for the code.

## Supplemental Figures

**Figure S1:**
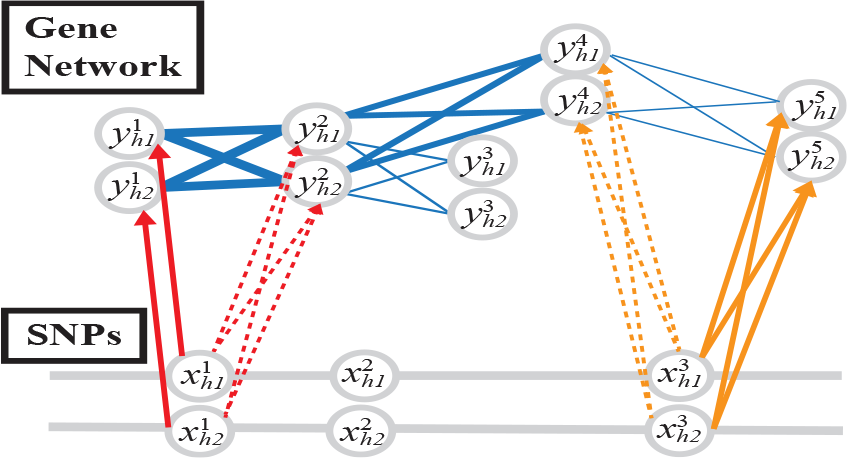
Illustration of inference in CiTruss. The estimated multi-level model contains eQTLs with direct effects (solid arrows). CiTruss infers eQTLs with indirect effects (dashed arrows) on the downstream genes in the gene network.

**Figure S2:**
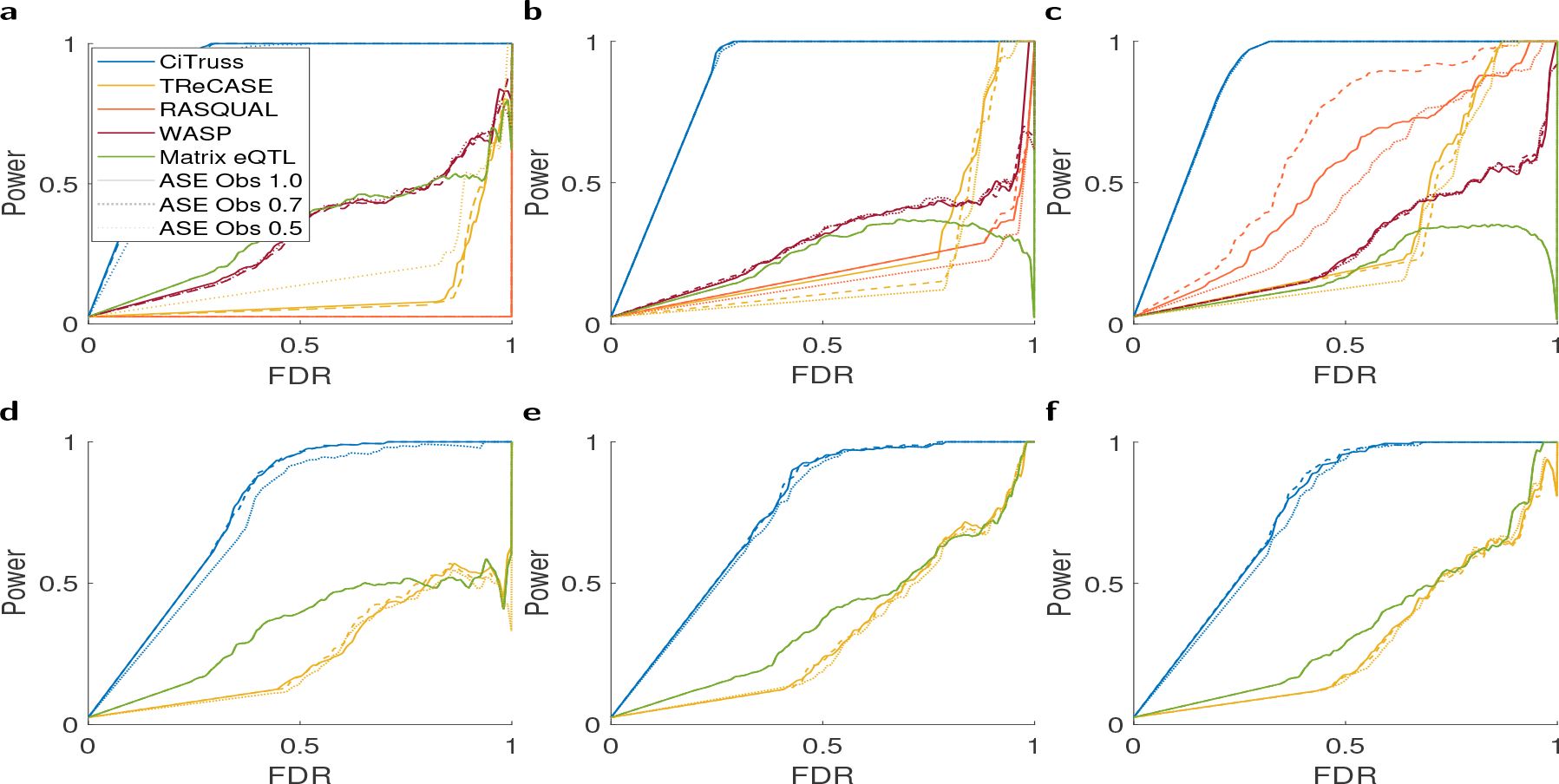
Comparison of CiTruss and other methods on the accuracy of *cis*-acting and *trans*-acting eQTLs in simulation. Accuracy of *cis*-acting eQTLs when the frequency of heterozygous genotypes is (a) 0.05, (b) 0.25, and (c) 0.45. Accuracy of *trans*-acting eQTLs when the frequency of heterozygous genotype is (d) 0.05, (e) 0.25, and (f) 0.45.

**Figure S3:**
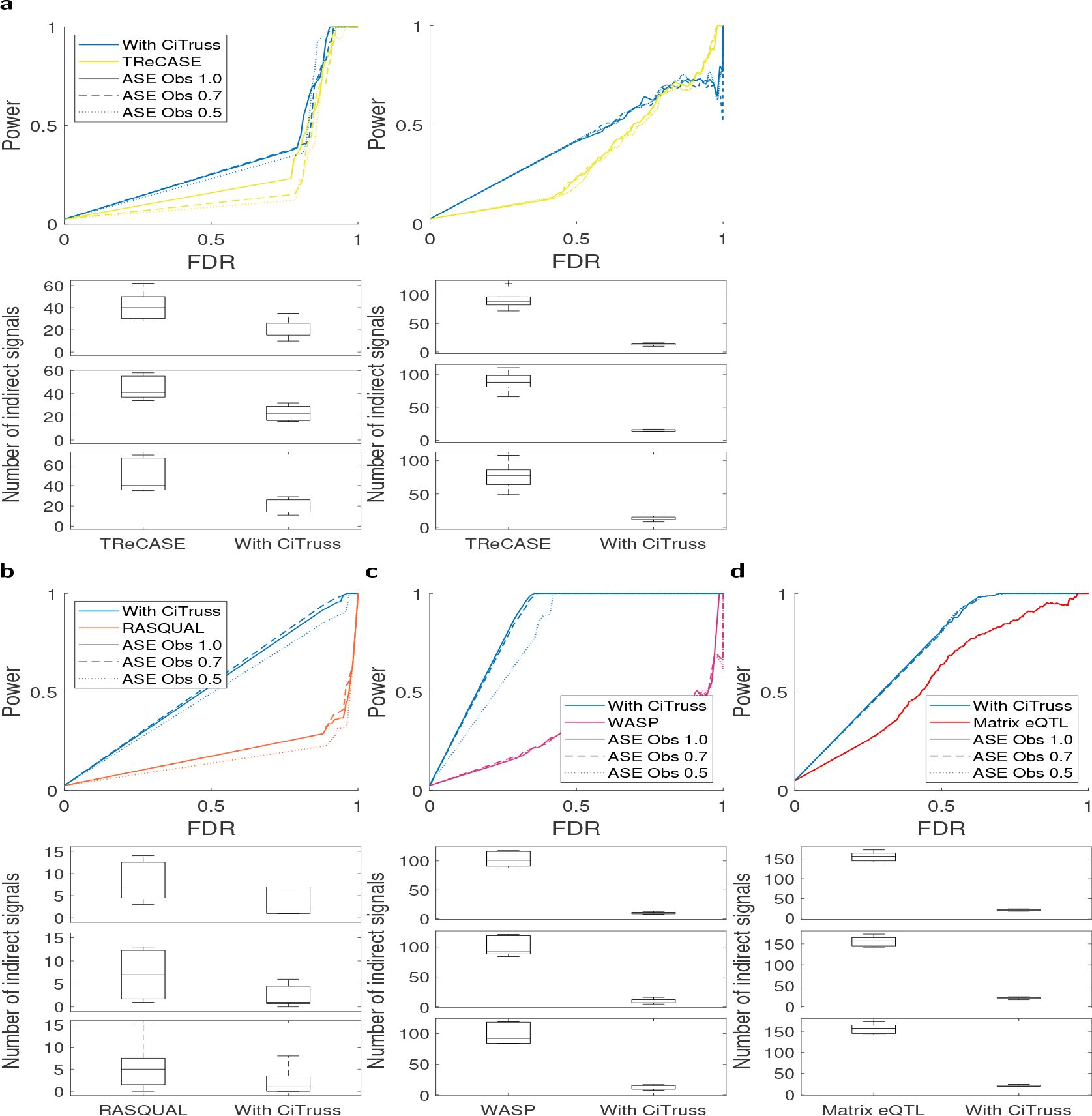
Performance of CiTruss partnered with the existing methods in simulation with the frequency of heterozygous genotypes 0.25. Statistically significant eQTLs (5% FDR) from TReCASE, RASQUAL, WASP, and Matrix eQTL were re-analyzed with CiTruss to select eQTLs with direct effects. The results from each of the existing methods were compared against those from re-analysis with CiTruss. (a) TReCASE on the accruacy of *cis*-acting eQTLs (left) and *trans*-acting eQTLs (right). Power at different FDR (top) and the number of true indirect eQTLs detected as eQTLs (bottom) are shown for datasets with the frequency of observed allele-specific expression levels 1.0, 0.7, and 0.5 from top to bottom. (b) RASQUAL on the accuracy of *cis*-acting eQTLs. (c) WASP on the accuracy of *cis*-acting eQTLs. (d) Matrix eQTL on the accuracy of eQTLs.

**Figure S4:**
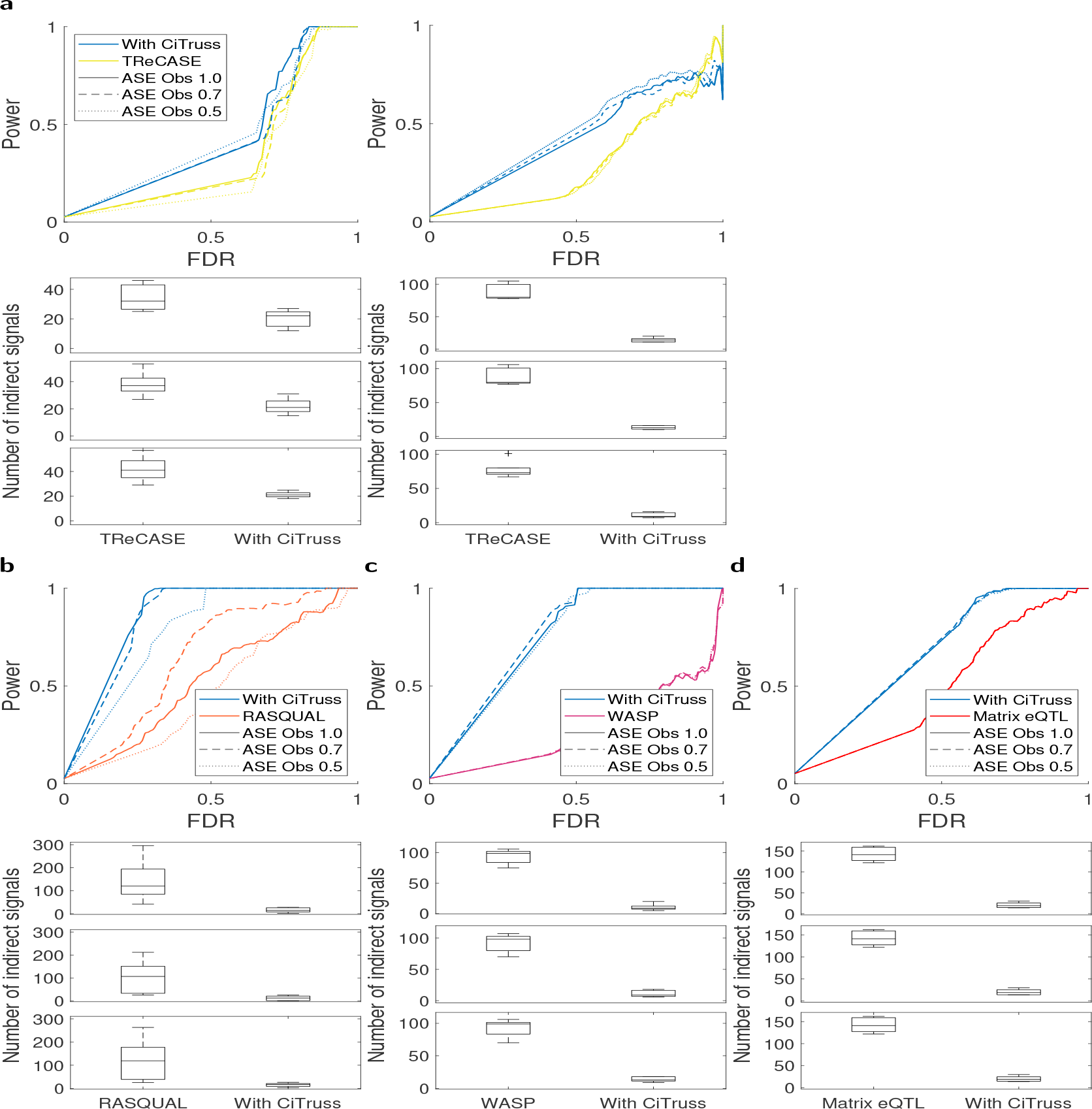
Performance of CiTruss partnered with the existing methods in simulation with frequency of heterozygous genotypes 0.45. Statistically significant eQTLs (5% FDR) from TReCASE, RASQUAL, WASP, and Matrix eQTLs were re-analyzed with CiTruss to select eQTLs with direct effects. The results from each of the existing methods were compared against those from re-analysis with CiTruss. (a) TReCASE on the accruacy of *cis*-acting eQTLs (left) and *trans*-acting eQTLs (right). Power at different FDR (top) and the number of true indirect eQTLs detected as eQTLs (bottom) are shown for datasets with the frequency of observed allele-specific expression levels 1.0, 0.7, and 0.5 from top to bottom. (b) RASQUAL on the accuracy of *cis*-acting eQTLs. (c) WASP on the accuracy of *cis*-acting eQTLs. (d) Matrix eQTL on the accuracy of eQTLs.

**Figure S5:**
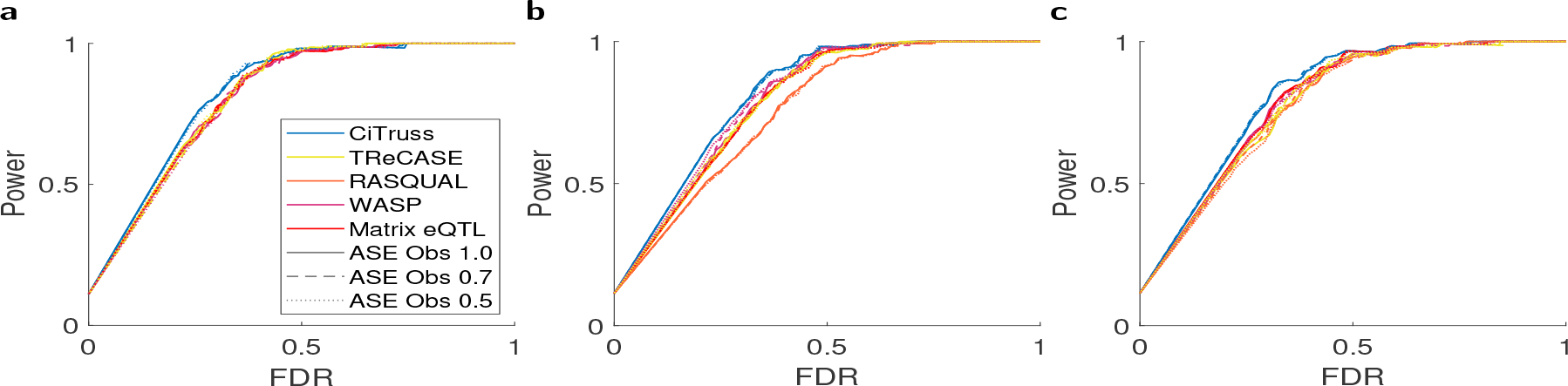
Accuracy of gene networks estimated by CiTruss in simulation. The gene networks estimated by CiTruss as a standalone method were compared with the networks from CiTruss re-analysis of statistically significant eQTLs (FDR 5%) from TReCASE, RASQUAL, WASP, and Matrix eQTL, when the frequency of heterozygous genotypes in samples is (a) 0.05, (b) 0.25, and (c) 0.45.

**Figure S6:**
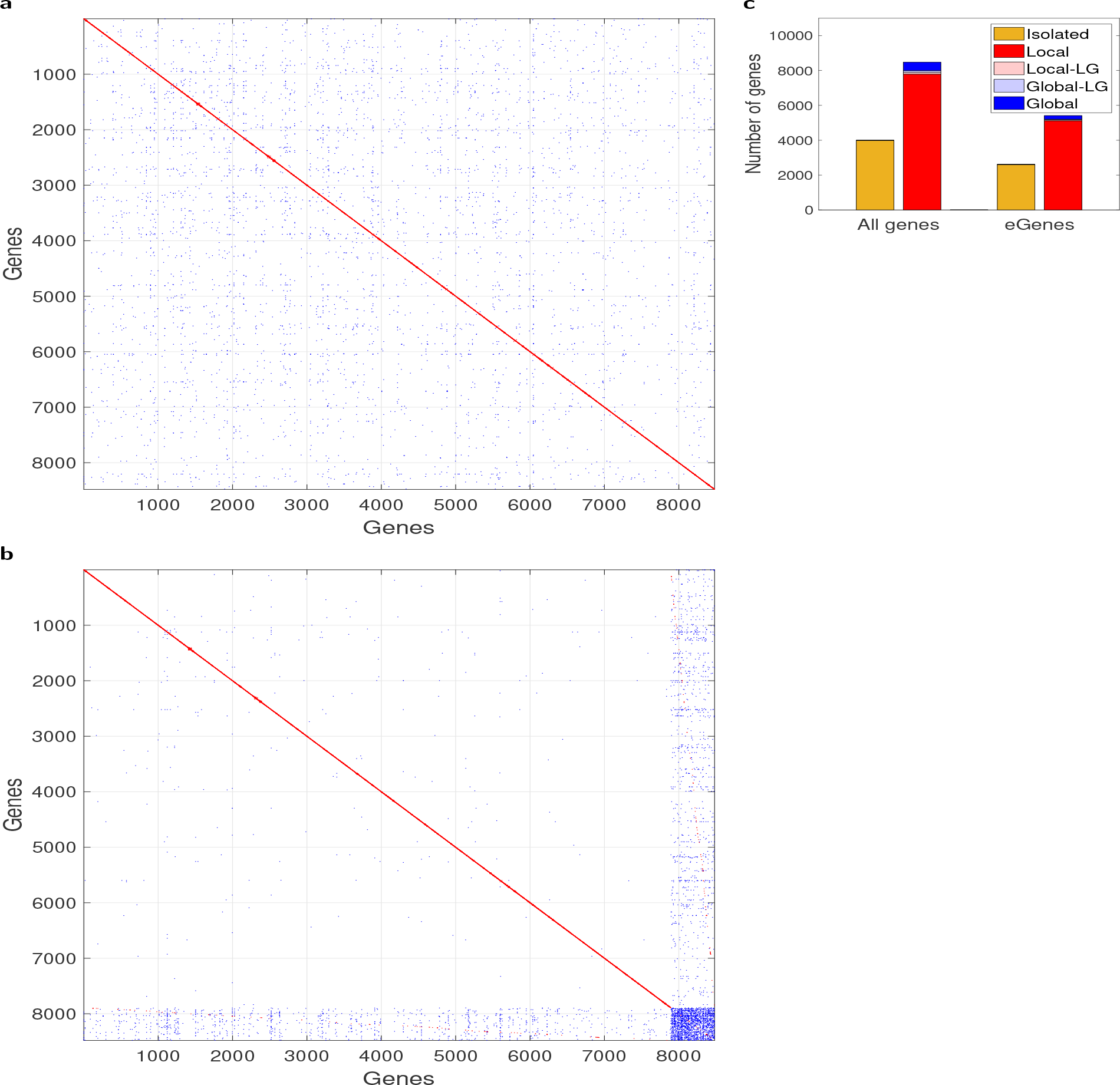
CiTruss gene network from whole blood GTEx data. The same results as in Figure 3 are shown for muscle skeletal.

**Figure S7:**
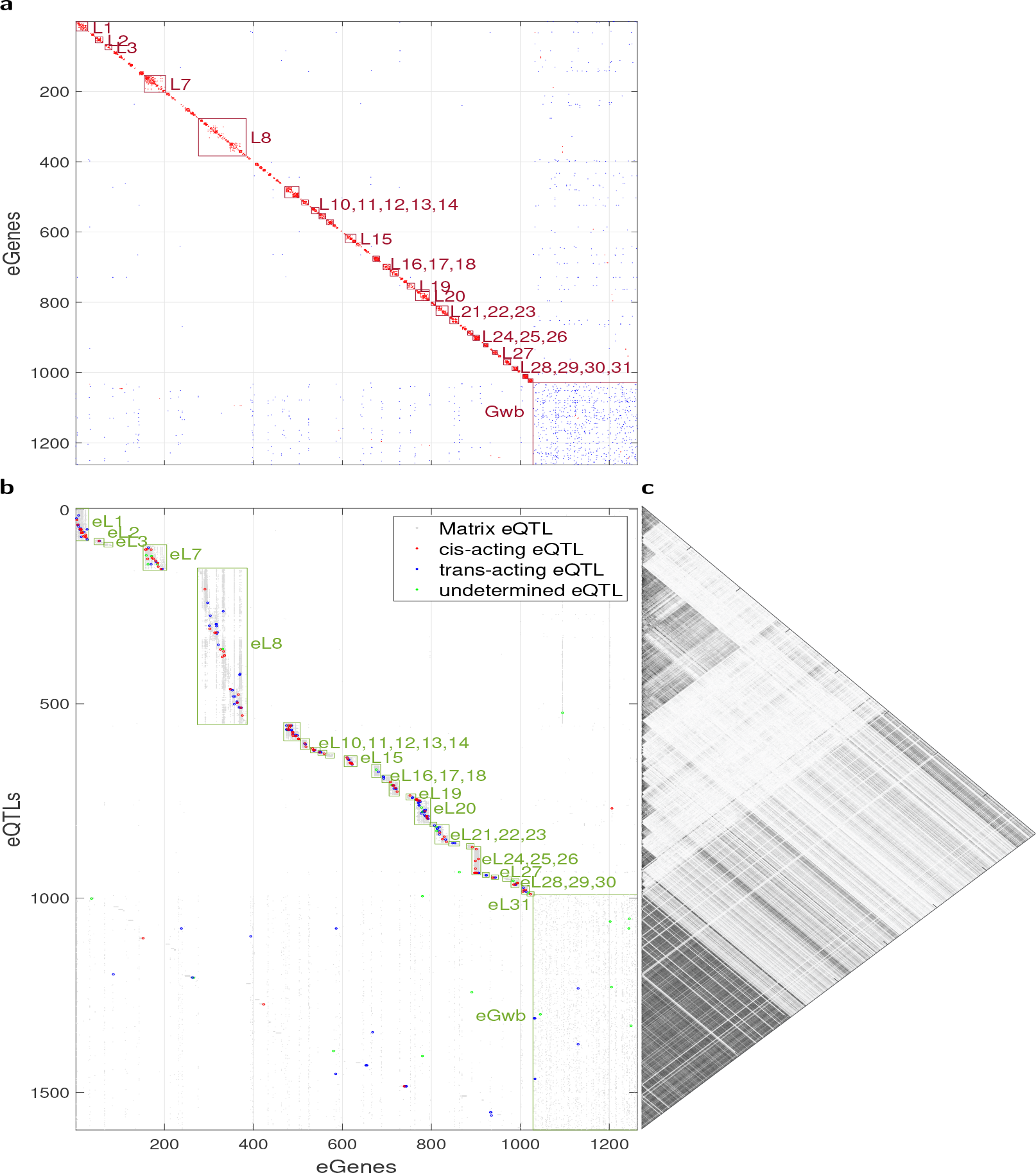
Matrix eQTL hotspots against CiTruss gene networks perturbed by eQTLs in muscle skeletal GTEx data. The same results as in Figure 4 are shown for muscle skeletal.

**Figure S8:**
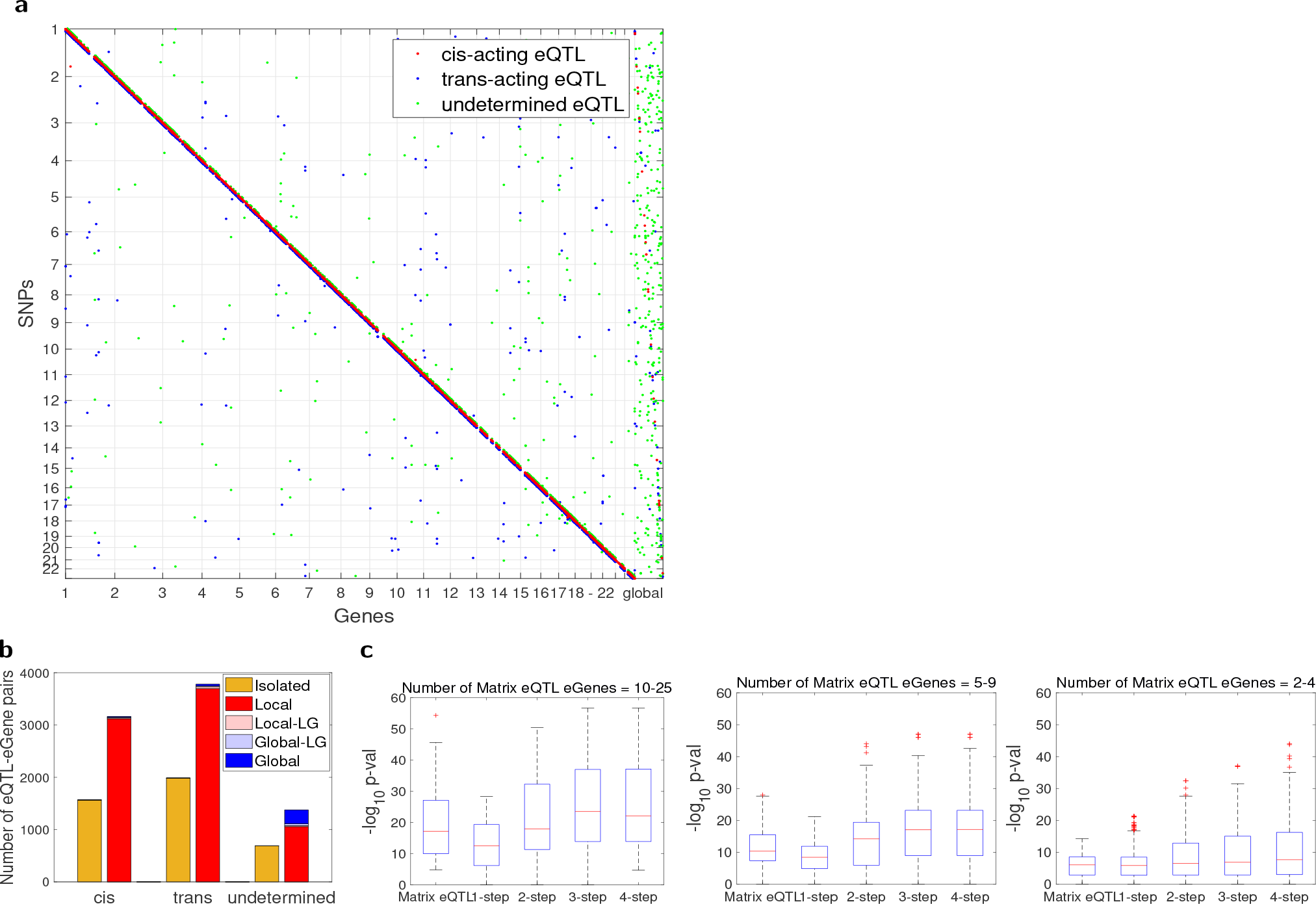
CiTruss eQTLs from whole blood GTEx data. The same results as in Figure 5 are shown for muscle skeletal.

**Figure S9:**
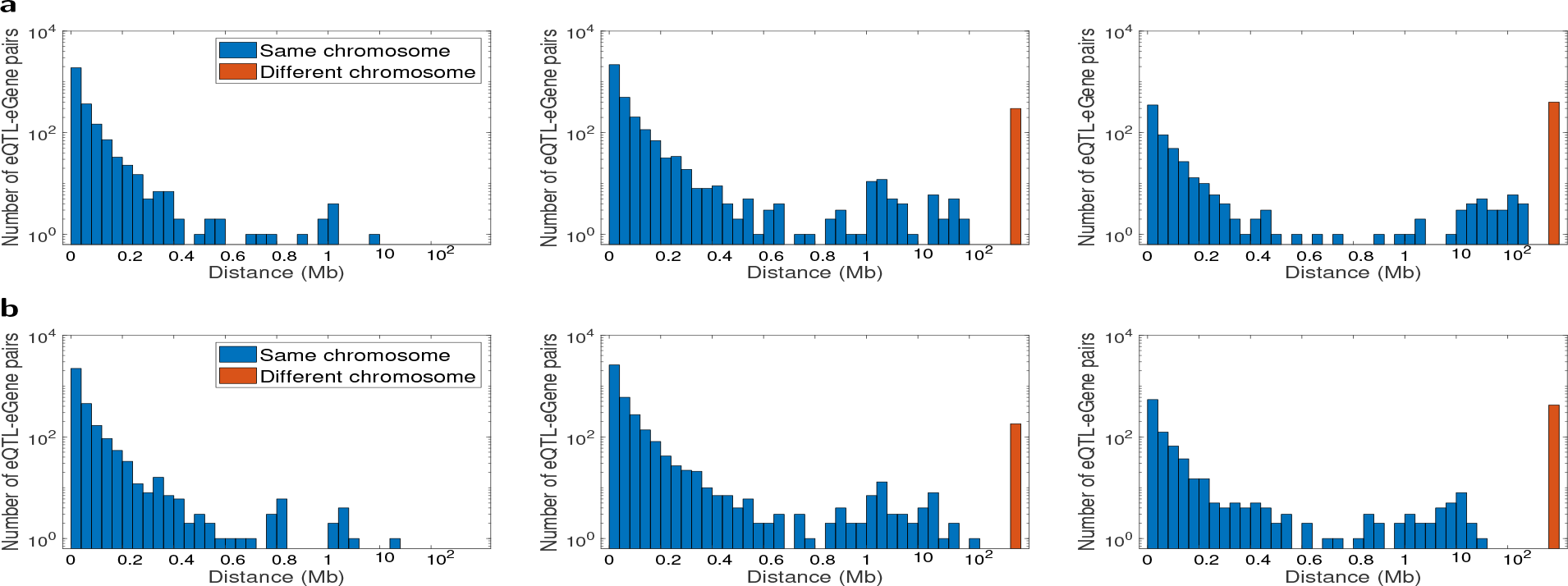
Distance between CiTruss eQTLs and their eGenes in GTEx data. (a) Whole blood and (b) muscle skeletal. *Cis*-acting eQTLs (left), *trans*-acting eQTLs (middle), and undetermined eQTLs (right).

**Figure S10:**
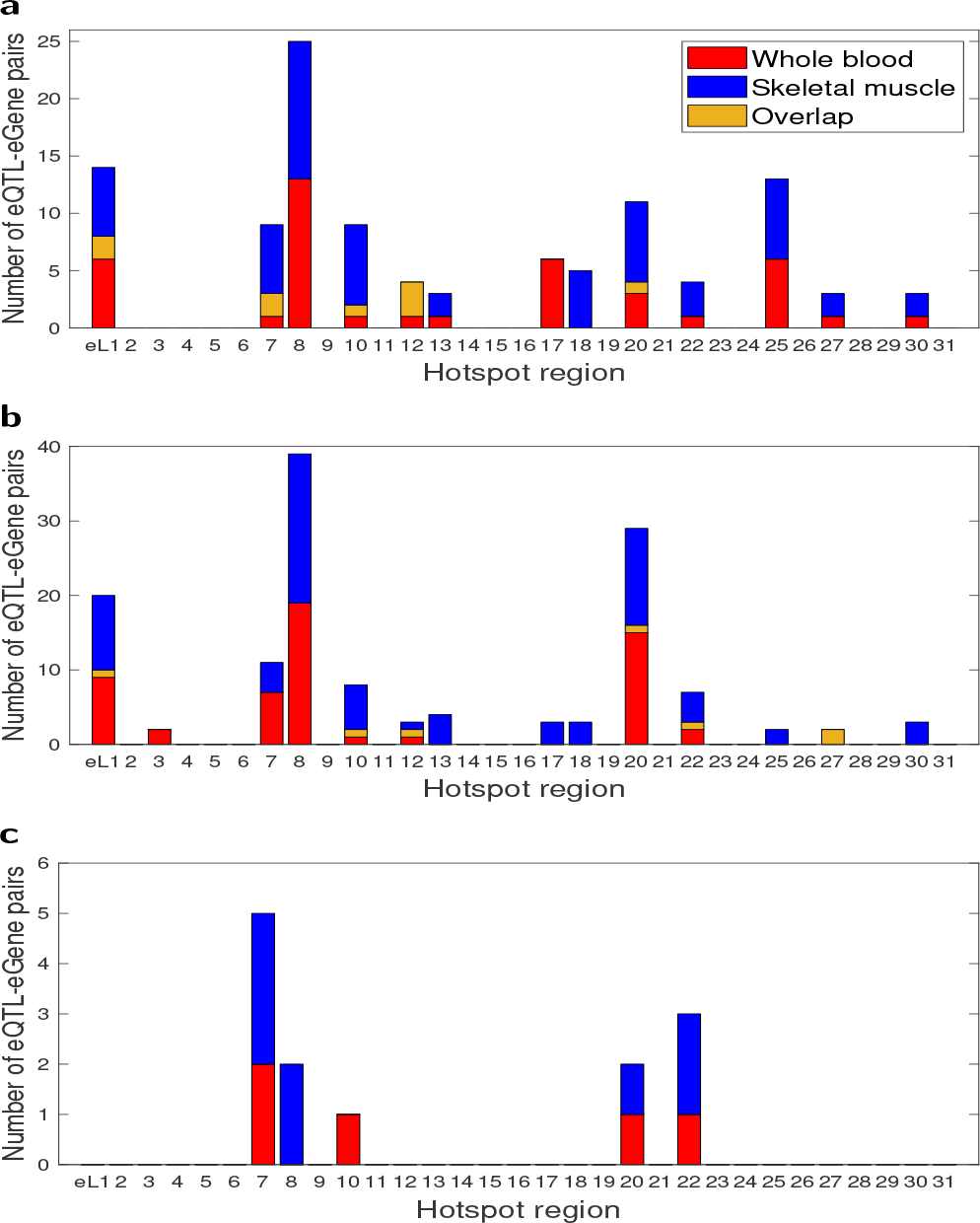
Comparison of CiTruss eQTLs for local subnetworks between whole blood and muscle skeletal in GTEx data. (a) *Cis*-acting eQTLs, (b) *trans*-acting eQTLs, and (c) undetermined eQTLs in whole blood, muscle skeletal, and the overlap between the two tissue types. The overlap shows the exact match in eQTLs between the two tissue types.

**Figure S11:**
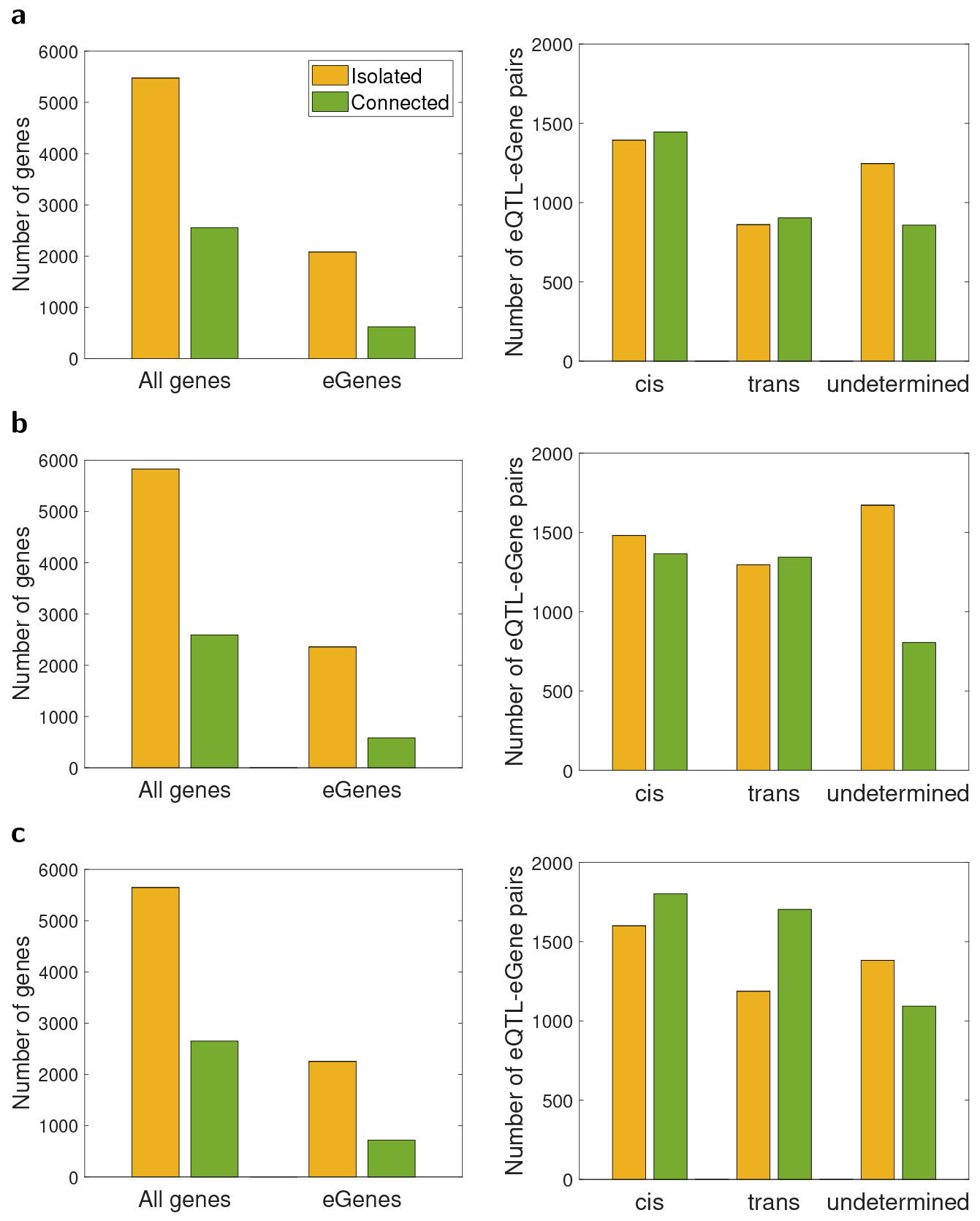
The composition of CiTruss gene networks and eQTLs from LG×SM AIL mouse data. (a) PFC, (b) STR, and (c) HIP. Genes and eGenes in gene networks (left) and eQTL-eGene pairs (right). Isolated: genes with no edges. Connected: genes with edges.

**Figure S12:**
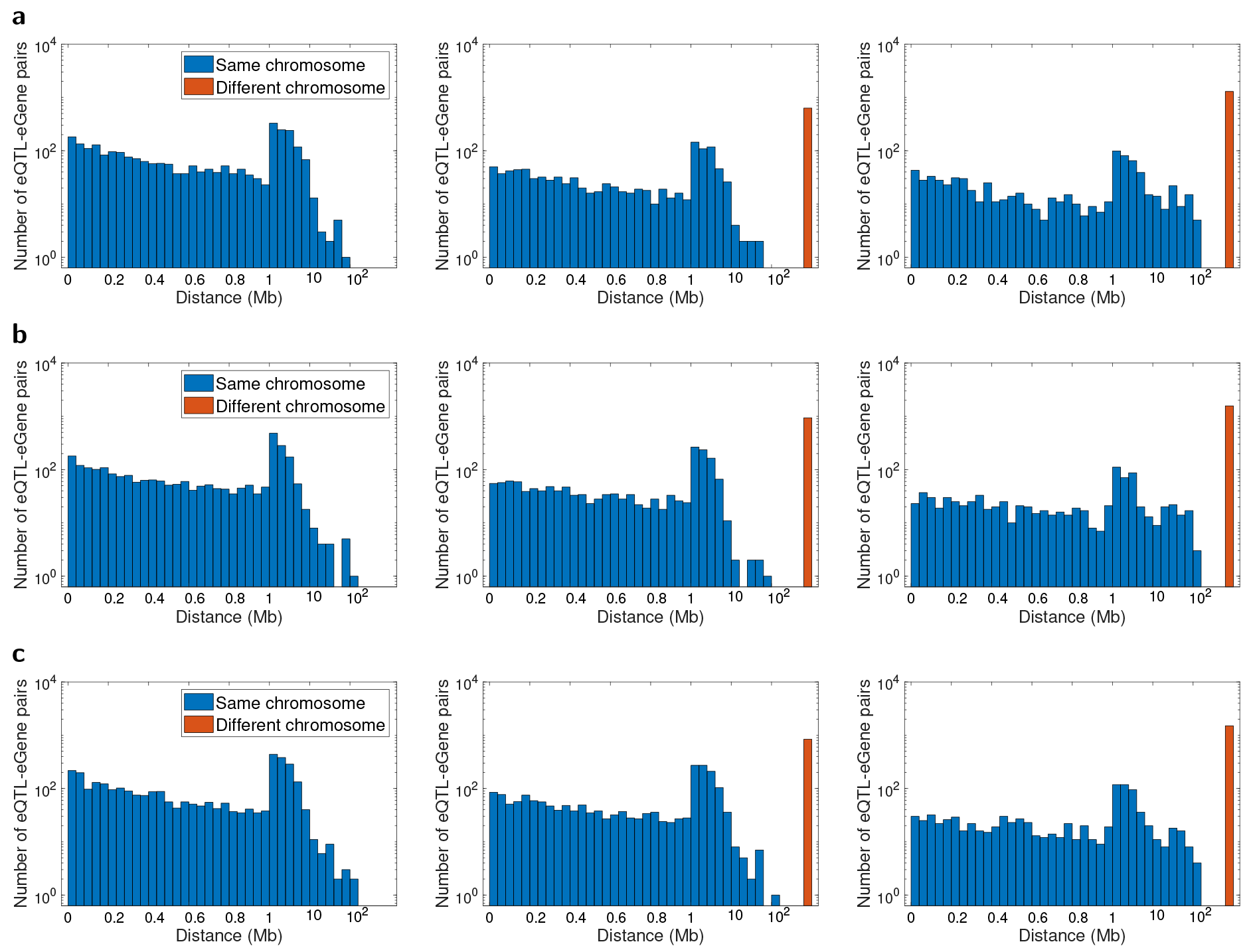
Distance between CiTruss eQTLs and their eGenes in LG×SM AIL mouse data. (a) PFC, (b) STR, and (c) HIP. *Cis*-acting eQTLs (left), *trans*-acting eQTLs (middle), and undetermined eQTLs (right).

**Figure S13:**
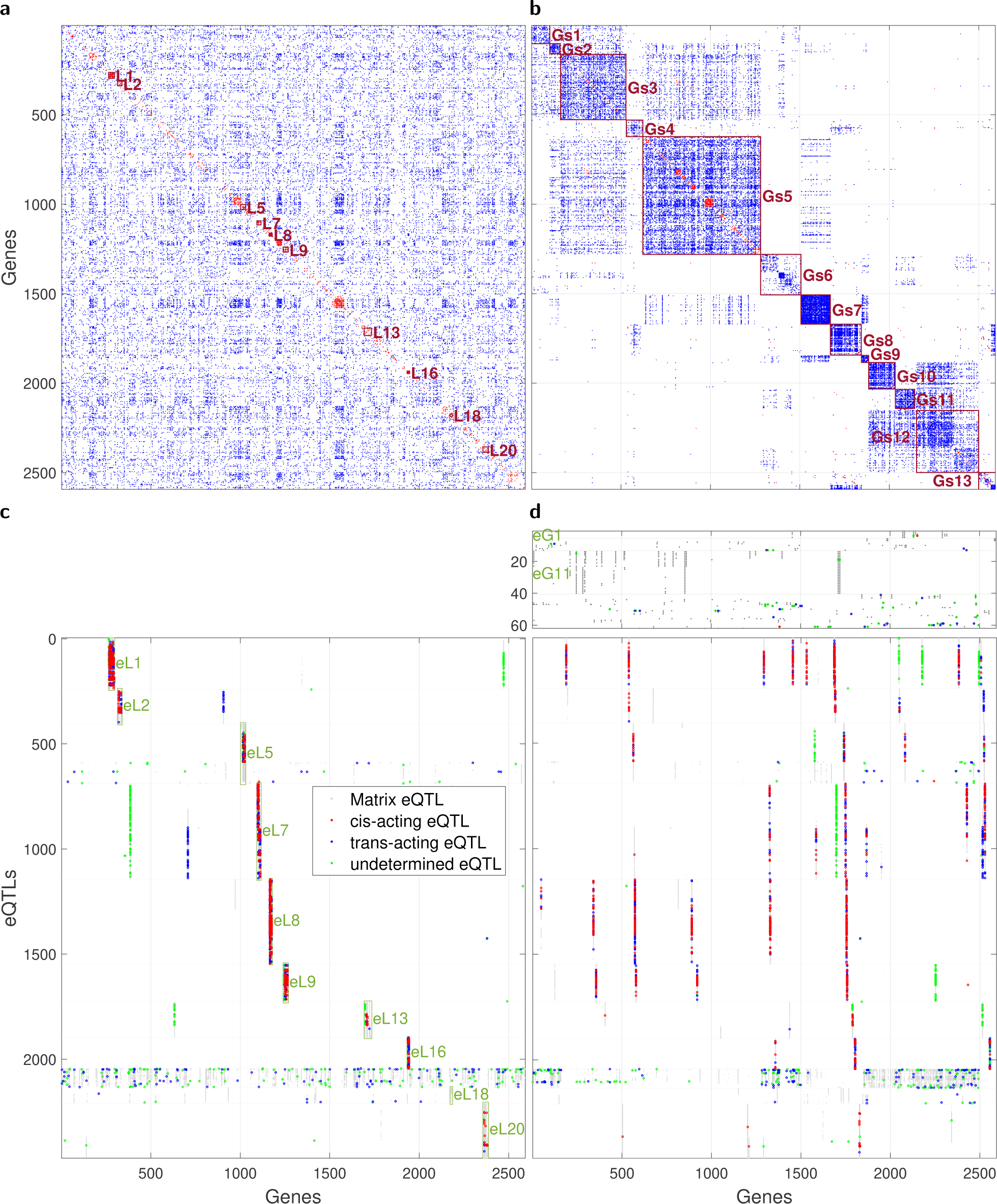
CiTruss gene network and eQTLs from LG×SM AIL mouse data for STR tissue. The same results as in Figure 6 are shown for STR.

**Figure S14:**
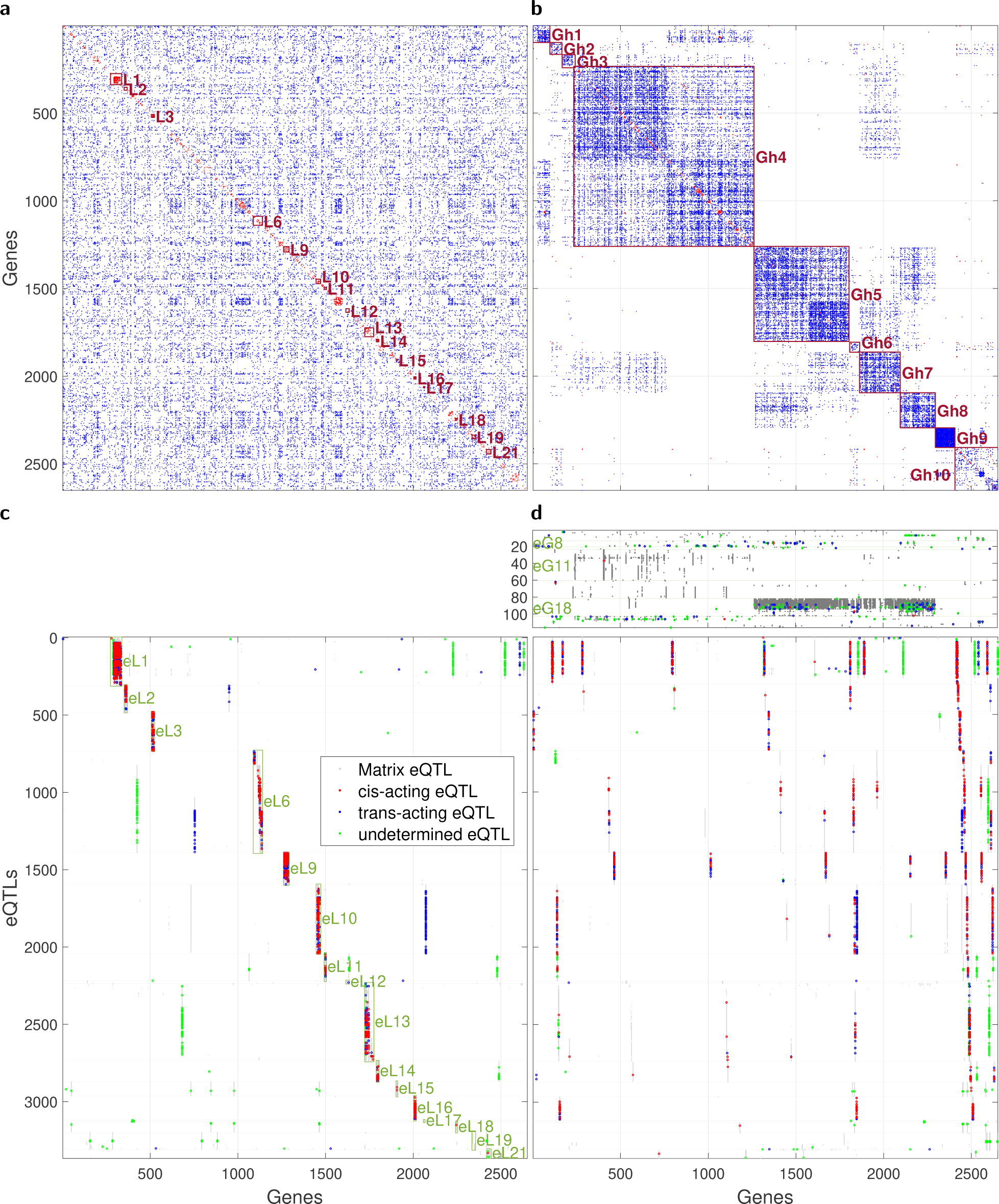
CiTruss gene network and eQTLs from LG×SM AIL mouse data for HIP tissue. The same results as in Figure 6 are shown for HIP.

**Figure S15:**
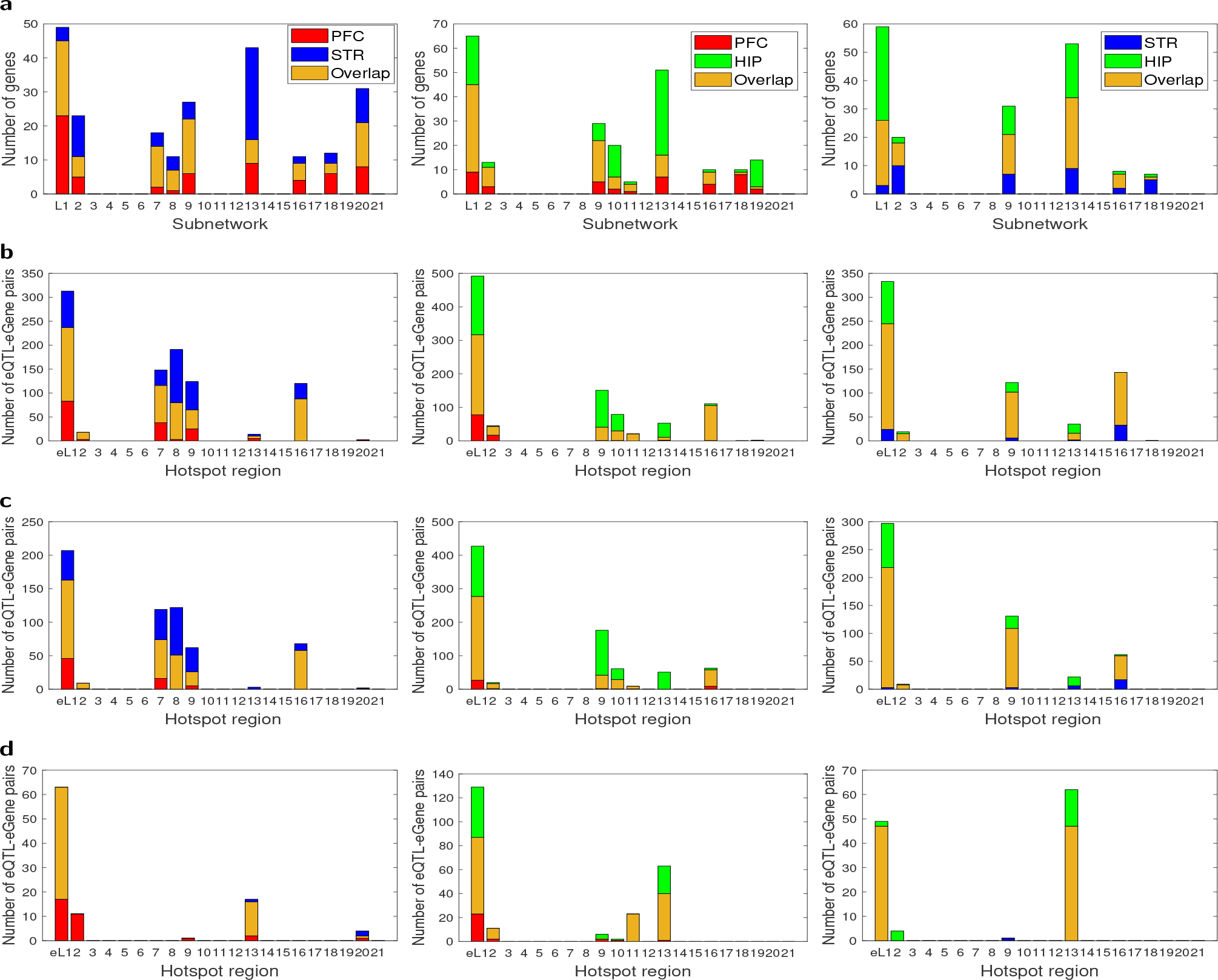
Comparison of CiTruss local subnetworks and eQTLs for local subnetworks across tissue types from LG SM AIL mouse data. (a) Local subnetworks, (b) *cis*-acting eQTLs, (c) *trans*-acting eQTLs, and (d) undetermined eQTLs. PFC vs STR (left), PFC vs HIP (middle), and STR vs HIP (right). The overlap in eQTLs shows the eQTLs within 500kb distance.

## Supplemental Tables

**Table S1:**
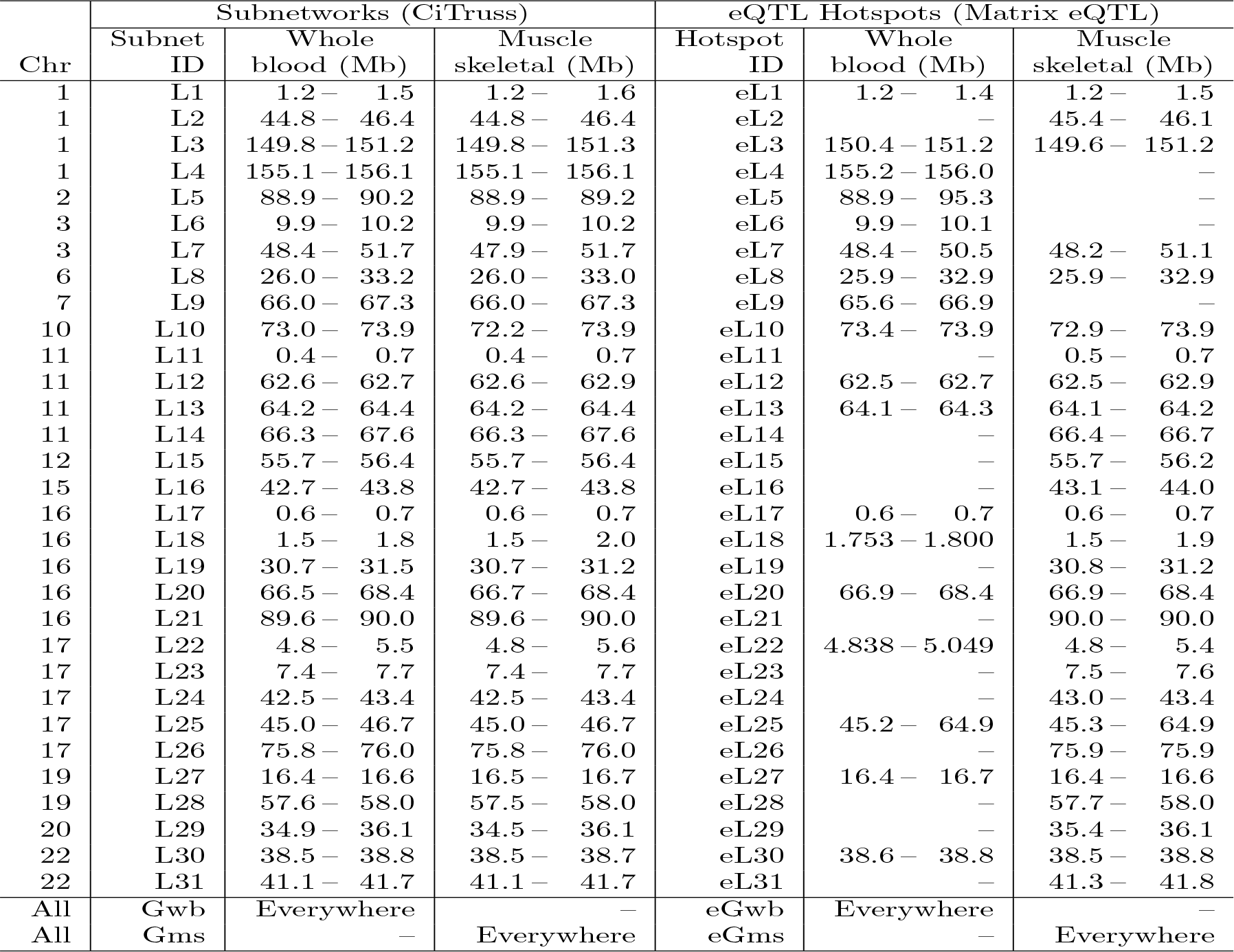
Genome regions for CiTruss subnetworks and Matrix eQTL hotspots in GTEx data.

**Table S2:** Genome regions of CiTruss local subnetworks and the corresponding Matrix eQTL hotspots in LG×SM AIL mouse data ^*^No edges in the local cluster, but edges to the global cluster.

| Chr | Local Subnetworks (CiTruss) |  |  |  | eQTL Hotspots (Matrix eQTL) |  |  |  |
| --- | --- | --- | --- | --- | --- | --- | --- | --- |
|  | Subnet ID | PFC (Mb) | STR (Mb) | HIP (Mb) | Hotspot ID | PFC (Mb) | STR (Mb) | HIP (Mb) |
| 2 | L1 | 113.8 – 129.1 | 118.5 – 127.0 | 102.3 – 129.1 | eL1 | 103.5 – 128.9 | 114.8 – 127.2 | 101.9 – 128.9 |
| 2 | L2 | 154.7 – 156.4* | 155.3 – 158.6 | 155.3 – 156.6* | eL2 | 148.9 – 158.6 | 154.3 – 157.6 | 154.3 – 158.6 |
| 3 | L3 | – | – | 122.0 – 127.8 | eL3 | – | – | 118.8 – 128.5 |
| 5 | L4 | 144.7 – 147.0 | – | – | eL4 | 144.3 – 147.2 | – | – |
| 7 | L5 | – | 28.5 – 34.4 | – | eL5 | – | 30.1 – 36.3 | – |
| 7 | L6 | – | – | 81.6 – 114.7* | eL6 | – | – | 81.9 – 115.0 |
| 7 | L7 | 109.5 – 114.7 | 105.7 – 114.7 | – | eL7 | 105.7 – 115.0 | 105.7 – 115.0 | – |
| 8 | L8 | 11.3 – 13.7 | 11.5 – 14.0 | – | eL8 | 11.2 – 12.3 | 11.2 – 14.5 | – |
| 8 | L9 | 82.9 – 85.7 | 83.3 – 85.1 | 82.9 – 85.8 | eL9 | 79.9 – 86.9 | 79.9 – 86.9 | 79.9 – 87.9 |
| 9 | L10 | 95.6 – 97.1 | – | 89.7 – 101.0 | eL10 | 93.2 – 97.5 | – | 83.9 – 100.2 |
| 9 | L11 | 119.3 – 120.6 | – | 119.3 – 120.6* | eL11 | 119.1 – 121.3 | – | 119.1 – 124.2 |
| 10 | L12 | – | – | 127.0 – 128.5* | eL12 | – | – | 127.2 – 129.9 |
| 11 | L13 | 68.9 – 70.7 | 68.9 – 78.3* | 68.9 – 78.3 | eL13 | 64.7 – 76.3 | 68.1 – 76.3 | 58.9 – 79.9 |
| 11 | L14 | – | – | 94.0 – 95.0 | eL14 | – | – | 92.3 – 96.5 |
| 12 | L15 | – | – | 17.9 – 17.9 | eL15 | – | – | 16.0 – 25.0 |
| 13 | L16 | 23.4 – 25.3 | 24.0 – 25.3 | 24.8 – 25.3 | eL16 | 20.2 – 29.6 | 22.8 – 29.6 | 20.2 – 29.6 |
| 13 | L17 | – | – | 103.8 – 103.9 | eL17 | – | – | 103.4 – 104.2 |
| 15 | L18 | 81.7 – 83.5* | 82.0 – 83.9* | 81.9 – 82.0* | eL18 | 81.9 – 84.4 | 82.0 – 86.6 | 81.7 – 85.4 |
| 16 | L19 | 31.3 – 32.0* | – | 32.0 – 38.2 | eL19 | 29.6 – 33.4 | – | 29.6 – 47.9 |
| 17 | L20 | 32.8 – 41.0 | 32.8 – 35.8 | – | eL20 | 9.2 – 43.0 | 30.8 – 43.0 | – |
| 17 | L21 | – | – | 32.8 – 35.8 | eL21 | – | – | 32.4 – 35.5 |

\*No edges in the local cluster, but edges to the global cluster.

**Table S3:** Genome regions of Matrix eQTL hotspots for CiTruss global subnetworks in LG×SM AIL mouse data.

| Chr | Hotspot ID | PFC (Mb) | STR (Mb) | HIP (Mb) |
| --- | --- | --- | --- | --- |
| 1 | eG1 | – | 71.0 – 71.1 | – |
| 1 | eG2 | – | – | 78.5 – 79.9 |
| 1 | eG3 | – | – | 175.6 – 178.8 |
| 4 | eG4 | – | 36.2 – 36.3 | – |
| 4 | eG5 | 40.3 – 41.7 | – | – |
| 4 | eG6 | 53.2 – 53.4 | – | – |
| 4 | eG7 | – | – | 155.3 – 155.3 |
| 5 | eG8 | 113.2 – 114.1 | – | 90.3 – 114.4 |
| 5 | eG9 | – | 125.0 – 125.1 | – |
| 6 | eG10 | – | 28.6 – 32.3 | – |
| 6 | eG11 | 127.2 – 128.3 | 124.8 – 128.1 | 123.6 – 133.9 |
| 6 | eG12 | 144.9 – 147.5 | – | – |
| 8 | eG13 | – | – | 11.6 – 14.0 |
| 9 | eG14 | – | 31.3 – 36.5 | – |
| 10 | eG15 | 115.5 – 115.5 | – | – |
| 10 | eG16 | – | – | 117.6 – 119.0 |
| 10 | eG17 | – | – | 121.6 – 122.1 |
| 11 | eG18 | 58.7 – 58.8 | – | 56.8 – 58.9 |
| 11 | eG19 | – | – | 105.2 – 106.8 |
| 12 | eG20 | 100.6 – 100.8 | – | – |
| 12 | eG21 | – | 107.5 – 107.7 | – |
| 13 | eG22 | – | – | 99.1 – 99.1 |
| 14 | eG23 | 54.4 – 54.5 | – | – |
| 17 | eG24 | – | – | 28.9 – 29.3 |
| 18 | eG25 | – | 23.5 – 23.5 | – |
| 18 | eG26 | – | 57.7 – 57.8 | – |
| 19 | eG27 | – | 22.7 – 23.3 | – |

## References

[1] Ritsert C Jansen and Jan-Peter Nap. Genetical genomics: the added value from segregation. Trends in Genetics, 17(7):388–391, 2001.

[2] Yang Li, Rainer Breitling, and Ritsert C Jansen. Generalizing genetical genomics: getting added value from environmental perturbation. Trends in Genetics, 24(10):518–524, 2008.

[3] Rainer Breitling, Yang Li, Bruno M Tesson, Jingyuan Fu, Chunlei Wu, Tim Wiltshire, Alice Gerrits, Leonid V Bystrykh, Gerald De Haan, Andrew I Su, et al. Genetical genomics: spotlight on QTL hotspots. PLoS Genetics, 4(10):e1000232, 2008.

[4] Wei Sun. A statistical framework for eQTL mapping using RNA-seq data. Biometrics, 68(1):1–11, 2012.

[5] Vasyl Zhabotynsky, Licai Huang, Paul Little, Yi-Juan Hu, Fernando Pardo-Manuel de Villena, Fei Zou, and Wei Sun. eQTL mapping using allele-specific count data is computationally feasible, powerful, and provides individual-specific estimates of genetic effects. PLoS Genetics, 18(3):e1010076, 2022.

[6] Natsuhiko Kumasaka, Andrew J. Knights, and Daniel J. Gaffney. Fine-mapping cellular QTLs with RASQUAL and ATAC-seq. Nature Genetics, 48(2):206–213, Feb 2016.

[7] Bryce van de Geijn, Graham McVicker, Yoav Gilad, and Jonathan K. Pritchard. WASP: allele-specific software for robust molecular quantitative trait locus discovery. Nature Methods, 12(11):1061–1063, 2015.

[8] Daphne Koller and Nir Friedman. Probabilistic graphical models: principles and techniques. MIT press, 2009.

[9] Lingxue Zhang and Seyoung Kim. Learning gene networks under SNP perturbations using eQTL datasets. PLoS Computational Biology, 10(2):e1003420, 2014.

[10] Calvin McCarter, Judie Howrylak, and Seyoung Kim. Learning gene networks underlying clinical phenotypes using SNP perturbation. PLoS Computational Biology, 16(10):e1007940, 2020.

[11] Andrey A. Shabalin. Matrix eQTL: ultra fast eQTL analysis via large matrix operations. Bioinformatics, 28(10):1353–1358, 2012.

[12] Vasyl Zhabotynsky, Licai Huang, Paul Little, Yi-Juan Hu, Fernando Pardo-Manuel de Villena, Fei Zou, and Wei Sun. eQTL mapping using allele-specific count data is computationally feasible, powerful, and provides individual-specific estimates of genetic effects. PLoS Genetics, 18(3):1–18, 2022.

[13] The GTEx Consortium et al. The GTEx consortium atlas of genetic regulatory effects across human tissues. Science, 369(6509):1318–1330, 2020.

[14] Natalia M Gonzales, Jungkyun Seo, Ana I Hernandez Cordero, Celine L St Pierre, Jennifer S Gregory, Margaret G Distler, Mark Abney, Stefan Canzar, Arimantas Lionikas, and Abraham A Palmer. Genome wide association analysis in a mouse advanced intercross line. Nature Communications, 9(1):5162, 2018.

[15] Diogo M Ribeiro, Simone Rubinacci, Anna Ramisch, Robin J Hofmeister, Emmanouil T Dermitzakis, and Olivier Delaneau. The molecular basis, genetic control and pleiotropic effects of local gene co-expression. Nature Communications, 12(1):4842, 2021.

[16] Shuli Liu, Yahui Gao, Oriol Canela-Xandri, Sheng Wang, Ying Yu, Wentao Cai, Bingjie Li, Ruidong Xiang, Amanda J Chamberlain, Erola Pairo-Castineira, et al. A multi-tissue atlas of regulatory variants in cattle. Nature Genetics, 54(9):1438–1447, 2022.

[17] Leeyoung Park. Population-specific long-range linkage disequilibrium in the human genome and its influence on identifying common disease variants. Scientific Reports, 9(1):11380, 2019.

[18] Abdelmajid El Hou, Dominique Rocha, Eric Venot, Véronique Blanquet, and Romain Philippe. Long-range linkage disequilibrium in french beef cattle breeds. Genetics Selection Evolution, 53:1–14, 2021.

[19] Pankhuri Singhal, Yogasudha Veturi, Scott M Dudek, Anastasia Lucas, Alex Frase, Kristel van Steen, Steven J Schrodi, David Fasel, Chunhua Weng, Rion Pendergrass, et al. Evidence of epistasis in regions of long-range linkage disequilibrium across five complex diseases in the UK Biobank and eMERGE datasets. The American Journal of Human Genetics, 110(4):575–591, 2023.

[20] Tianshun Gao and Jiang Qian. EnhancerAtlas 2.0: an updated resource with enhancer annotation in 586 tissue/cell types across nine species. Nucleic Acids Research, 48(D1):D58–D64, 2019.

[21] Qiong Zhang, Wei Liu, Hong-Mei Zhang, Gui-Yan Xie, Ya-Ru Miao, Mengxuan Xia, and An-Yuan Guo. hTFtarget: A comprehensive database for regulations of human transcription factors and their targets. Genomics, Proteomics & Bioinformatics, 18(2):120–128, 2020.

[22] Jaime A Castro-Mondragon, Rafael Riudavets-Puig, Ieva Rauluseviciute, Roza Berhanu Lemma, Laura Turchi, Romain Blanc-Mathieu, Jeremy Lucas, Paul Boddie, Aziz Khan, Nicolás Manosalva Pérez, Oriol Fornes, Tiffany Y Leung, Alejandro Aguirre, Fayrouz Hammal, Daniel Schmelter, Damir Baranasic, Benoit Ballester, Albin Sandelin, Boris Lenhard, Klaas Vandepoele, Wyeth W Wasserman, François Parcy, and Anthony Mathelier. JASPAR 2022: the 9th release of the open-access database of transcription factor binding profiles. Nucleic Acids Research, 50(D1):D165–D173, 2021.

[23] Andrew D. Rouillard, Gregory W. Gundersen, Nicolas F. Fernandez, Zichen Wang, Caroline D. Monteiro, Michael G. McDermott, and Avi Ma’ayan. The harmonizome: a collection of processed datasets gathered to serve and mine knowledge about genes and proteins. Database, 2016, 2016.

[24] Orsolya Liska, Balázs Bohár, András Hidas, Tamás Korcsmáros, Balázs Papp, Dávid Fazekas, and Eszter Ari. TFLink: an integrated gateway to access transcription factor–target gene interactions for multiple species. Database, 2022, 2022.

[25] Daniel Munro, Tengfei Wang, Apurva S Chitre, Oksana Polesskaya, Nava Ehsan, Jianjun Gao, Alexander Gusev, Leah C Solberg Woods, Laura M Saba, Hao Chen, et al. The regulatory landscape of multiple brain regions in outbred heterogeneous stock rats. Nucleic Acids Research, 50(19):10882–10895, 2022.

[26] Mattia Forcato, Chiara Nicoletti, Koustav Pal, Carmen Maria Livi, Francesco Ferrari, and Silvio Bicciato. Comparison of computational methods for Hi-C data analysis. Nature Methods, 14(7):679–685, 2017.

[27] Xiang Zhou and Matthew Stephens. Genome-wide efficient mixed-model analysis for association studies. Nature Genetics, 44(7):821–824, 2012.

[28] Maryam Onifade, Marie-Hélène Roy-Gagnon, Marie-Élise Parent, and Kelly M Burkett. Comparison of mixed model based approaches for correcting for population substructure with application to extreme phenotype sampling. BMC Genomics, 23(1):98, 2022.

[29] Jun Ho Yoon and Seyoung Kim. EiGLasso for scalable sparse Kronecker-sum inverse covariance estimation. The Journal of Machine Learning Research, 23(1):4733–4771, 2022.

[30] Atray Dixit, Oren Parnas, Biyu Li, Jenny Chen, Charles P Fulco, Livnat Jerby-Arnon, Nemanja D Marjanovic, Danielle Dionne, Tyler Burks, Raktima Raychowdhury, et al. Perturb-Seq: dissecting molecular circuits with scalable single-cell RNA profiling of pooled genetic screens. Cell, 167(7):1853–1866, 2016.

[31] Daniel Schraivogel, Andreas R Gschwind, Jennifer H Milbank, Daniel R Leonce, Petra Jakob, Lukas Mathur, Jan O Korbel, Christoph A Merten, Lars Velten, and Lars M Steinmetz. Targeted Perturb-Seq enables genome-scale genetic screens in single cells. Nature Methods, 17(6):629–635, 2020.

[32] Calvin McCarter and Seyoung Kim. Large-scale optimization algorithms for sparse conditional Gaussian graphical models. In Proceedings of the 19th International Conference on Artificial Intelligence and Statistics, volume 51, pages 528–537. PMLR, 2016.

[33] Shaun Purcell, Benjamin Neale, Kathe Todd-Brown, Lori Thomas, Manuel A R Ferreira, David Bender, Julian Maller, Pamela Sklar, Paul I W de Bakker, Mark J Daly, and Pak C Sham. PLINK: a tool set for whole-genome association and population-based linkage analyses. The American Journal of Human Genetics, 81(3):559–575, 2007.

[34] Petr Danecek, James K Bonfield, Jennifer Liddle, John Marshall, Valeriu Ohan, Martin O Pollard, Andrew Whitwham, Thomas Keane, Shane A McCarthy, Robert M Davies, and Heng Li. Twelve years of SAMtools and BCFtools. GigaScience, 10(2), 2021.

[35] Brian L. Browning and Sharon R. Browning. Genotype imputation with millions of reference samples. The American Journal of Human Genetics, 98(1):116–126, 2016.

[36] Sharon R. Browning and Brian L. Browning. Rapid and accurate haplotype phasing and missing-data inference for whole-genome association studies by use of localized haplotype clustering. The American Journal of Human Genetics, 81(5):1084–1097, 2007.

[37] Tianshun Gao and Jiang Qian. EAGLE: An algorithm that utilizes a small number of genomic features to predict tissue/cell type-specific enhancer-gene interactions. PLoS Computational Biology, 15(10):1–22, 2019.

[38] Thomas Nall Eden Greville. Note on the generalized inverse of a matrix product. SIAM Review, 8(4): 518–521, 1966.

[39] Charis L Himeda, Jeffrey A Ranish, and Stephen D Hauschka. Quantitative proteomic identification of MAZ as a transcriptional regulator of muscle-specific genes in skeletal and cardiac myocytes. Molecular and cellular biology, 28(20):6521–6535, 2008.

[40] Christian Schutt, Alix Hallmann, Salma Hachim, Ina Klockner, Melissa Valussi, Ann Atzberger, Johannes Graumann, Thomas Braun, and Thomas Boettger. Linc-MYH configures INO80 to regulate muscle stem cell numbers and skeletal muscle hypertrophy. The EMBO Journal, 39(22):e105098, 2020.

[41] C Papadopoulos, M Svingou, K Kekou, S Vergnaud, S Xirou, G Niotakis, and GK Papadimas. Aldolase a deficiency: Report of new cases and literature review. Molecular Genetics and Metabolism Reports, 27: 100730, 2021.

[42] YiLi Wang, WonHee Han, SeokMin Yun, and JinKwan Han. Identification of protein phosphatase 4 catalytic subunit as a Wnt promoting factor in pan-cancer and xenopus early embryogenesis. Scientific Reports, 13(1):10240, 2023.

[43] Jessica X Chong, Jared C Talbot, Emily M Teets, Samantha Previs, Brit L Martin, Kathryn M Shively, Colby T Marvin, Arthur S Aylsworth, Reem Saadeh-Haddad, Ulrich A Schatz, et al. Mutations in MYLPF cause a novel segmental amyoplasia that manifests as distal arthrogryposis. The American Journal of Human Genetics, 107(2):293–310, 2020.

[44] MDE Deato and R Tjian. An unexpected role of TAFs and TRFs in skeletal muscle differentiation: switching core promoter complexes. In Cold Spring Harbor symposia on quantitative biology, volume 73, pages 217–225. Cold Spring Harbor Laboratory Press, 2008.

[45] Haiying Zhou, Bo Wan, Ivan Grubisic, Tommy Kaplan, and Robert Tjian. TAF7L modulates brown adipose tissue formation. Elife, 3:e02811, 2014.

[46] Fengyuan Chen, Jiajian Zhou, Yuying Li, Yu Zhao, Jie Yuan, Yang Cao, Lijun Wang, Zongkang Zhang, Baoting Zhang, Chi Chiu Wang, et al. YY1 regulates skeletal muscle regeneration through controlling metabolic reprogramming of satellite cells. The EMBO Journal, 38(10):e99727, 2019.

[47] Liang Zhou, Kun Sun, YU Zhao, Suyang Zhang, Xuecong Wang, Yuying Li, Leina Lu, Xiaona Chen, Fengyuan Chen, Xichen Bao, et al. Linc-YY1 promotes myogenic differentiation and muscle regeneration through an interaction with the transcription factor YY1. Nature Communications, 6(1):10026, 2015.

[48] Kun Sun, Leina Lu, Huating Wang, and Hao Sun. Genome-wide profiling of YY1 binding sites during skeletal myogenesis. Genomics Data, 2:89–91, 2014.

[49] Leina Lu, Liang Zhou, Eric Z Chen, Kun Sun, Peiyong Jiang, Lijun Wang, Xiaoxi Su, Hao Sun, and Huating Wang. A novel YY1-mir-1 regulatory circuit in skeletal myogenesis revealed by genome-wide prediction of YY1-miRNA network. PLoS One, 7(2):e27596, 2012.

[50] Casey L Sexton, Joshua S Godwin, Mason C McIntosh, Bradley A Ruple, Shelby C Osburn, Blake R Hollingsworth, Nicholas J Kontos, Philip J Agostinelli, Andreas N Kavazis, Tim N Ziegenfuss, et al. Skeletal muscle DNA methylation and mRNA responses to a bout of higher versus lower load resistance exercise in previously trained men. Cells, 12(2):263, 2023.

[51] Tanzila Mukhtar, Jeremie Breda, Alice Grison, Zahra Karimaddini, Pascal Grobecker, Dagmar Iber, Christian Beisel, Erik van Nimwegen, and Verdon Taylor. Tead transcription factors differentially regulate cortical development. Scientific Reports, 10(1):4625, 2020.

[52] Heba M El-Tahir, Mekky M Abouzied, Rainer Gallitzendoerfer, Volkmar Gieselmann, and Sebastian Franken. Hepatoma-derived growth factor-related protein-3 interacts with microtubules and promotes neurite outgrowth in mouse cortical neurons. Journal of Biological Chemistry, 284(17):11637–11651, 2009.

[53] Alisa Atkins, Michelle J Xu, Maggie Li, Nathaniel P Rogers, Marina V Pryzhkova, and Philip W Jordan. SMC5/6 is required for replication fork stability and faithful chromosome segregation during neurogenesis. Elife, 9:e61171, 2020.

[54] Tatiana Popovitchenko, Yongkyu Park, Nicholas F Page, Xiaobing Luo, Zeljka Krsnik, Yuan Liu, Iva Salamon, Jessica D Stephenson, Matthew L Kraushar, Nicole L Volk, et al. Translational derepression of Elavl4 isoforms at their alternative 5’ UTRs determines neuronal development. Nature Communications, 11(1):1674, 2020.

[55] Frank R Sharp and Myriam Bernaudin. HIF1 and oxygen sensing in the brain. Nature Reviews Neuroscience, 5(6):437–448, 2004.

[56] Hady Felfly, Alexander C Zambon, Jin Xue, Alysson Muotri, Dan Zhou, Evan Y Snyder, and Gabriel G Haddad. Severe hypoxia: consequences to neural stem cells and neurons. Journal of Neurology Research, 1(5), 2011.

[57] Helen T McKenna, Andrew J Murray, and Daniel S Martin. Human adaptation to hypoxia in critical illness. Journal of Applied Physiology, 129(4):656–663, 2020.

[58] Angeliki Karagiota, Ilias Mylonis, George Simos, and Georgia Chachami. Protein phosphatase PPP3CA (calcineurin A) down-regulates hypoxia-inducible factor transcriptional activity. Archives of biochemistry and biophysics, 664:174–182, 2019.

